# Investigating the Effect of GLU283 Protonation State on the Conformational Heterogeneity of CCR5 by Molecular Dynamics Simulations

**DOI:** 10.1101/2023.08.17.553662

**Authors:** Berna Dogan, Serdar Durdagi

## Abstract

CCR5 is one of the co-receptors for HIV-1 entry into host cells and is class A GPCR. This receptor has vital roles in the immune system and is involved in the pathogenesis of different diseases. Various studies were conducted to understand its activation mechanism including structural studies in which inactive and active states of the receptor were determined in complex with various binding partners. These determined structures provided opportunities to perform molecular dynamics simulations (MD) and analyze conformational changes observed in protein structures. The atomic level dynamical studies allow us to explore the effects of ionizable residues in the receptor. Here, our aim was to investigate the changes observed in the conformation of CCR5 when it is in complex with inhibitor maraviroc (MRV), an approved anti-HIV drug or HIV-1 envelope protein GP120 in comparison to when the receptor was in *apo* form. In our simulations, we considered both ionized and protonated states of ionizable binding site residue GLU283^7.39^ in CCR5 as the protonation state of this residue was considered ambiguously in previous studies. Our simulation results suggested that in fact, the change in the protonation state of GLU283^7.39^ caused interaction profiles to be different between CCR5 and its binding partners, GP120 or MRV. We observed that when the protonated state of GLU283^7.39^ was considered in complex with envelope protein GP120, there were substantial structural changes in CCR5 indicating it adopts more of an active-like conformation. On the other hand, CCR5 when it was in complex with MRV always adopted inactive conformation regardless of the protonation state. Hence, CCR5 coreceptor displays conformational heterogeneity not only based on its binding partner but also on the state of the protonation state of a binding site residue GLU283^7.39^. This outcome is also in accordance with some studies showing that GP120 binding could activate signaling pathways. Additionally, this outcome could also have critical implications for the discovery of novel CCR5 inhibitors to be used as anti-HIV drugs by in silico methods such as molecular docking since consideration of the protonated state of GLU283^7.39^ could be required.

## 1. INTRODUCTION

Chemokine receptors are a family of receptors that belong to class A G protein-coupled receptors (GPCRs) with crucial physiological roles specifically in the immune system.^1^ One well-studied member of this family C-C receptor type 5 (CCR5)^2^ is involved in the pathologies of various diseases including but not limited to inflammatory diseases, cancers, and infectious diseases.^3–4^ This receptor was also associated with the inflammatory complications of COVID-19.^5^ More intriguingly, CCR5 was identified as the principal coreceptor for M-tropic HIV-1 entry into the host cell.^6–7^ As such, this chemokine receptor is established as a well-known drug target and numerous research has been conducted to discover and develop molecules to inhibit its function. As a result, Maraviroc (MRV) was approved by the FDA to be used as a part of antiretroviral therapy for HIV infection in 2007.

CCR5 like other GPCRs is involved in cell signaling by binding to heteromeric G-proteins or β-arrestins.^8^ As other GPCRs, CCR5 cycles between two different states; active and inactive which differ in the orientation of their transmembrane (TM) helixes 5-6-7 specifically on intracellular sides. CCR5 is activated exogenously to initiate downstream signaling by four similar chemokines: MIP-1α, MIP-1β, RANTES, and MPC-2. This chemokine receptor adopts inactive conformation while bounded by antagonist or inverse agonists like MRV. Various studies were conducted to resolve the structure of CCR5 receptor with different ligands as well as in *apo* form. The first structure of CCR5 was resolved with MRV by X-ray crystallography in 2013 by Tan et al.^9^ and the receptor as expected was in inactive state. In 2018, in a crucial study Shaik et al. determined CCR5 structure with envelope protein of HIV-1,GP120 using cryo-electron microscopy (cryo-EM). In 2021, by two different studies and cryo-EM, active structures of CCR5 became available. Isikana et al. reported super agonist, [6P4]CCL5 bound structure^10^ and discussed possible activation mechanism for CCR5. Zhang et al. reported natural ligand bound structures of CCR5 in active form with chemokines RANTES and MIP-1α.^11^ They also reported CCR5 structure in *apo* form coupled with G_i_ protein. (see Table S1 in supporting information for details). All the structures determined were crucial in understanding the activation mechanism of CCR5.

CCR5 as the coreceptor of HIV, can engage with virus envelope glycoprotein GP120 after the virus binds to CD4 receptor of the host cell and eventually promotes the virus entry. GP120 was postulated to interact with two chemokine recognition sites on CCR5, same as chemokines and chemokine analogs. In fact, Shaik et al. validated this hypothesis with their cryo-EM structure of CC5 in complex with HIV-1 GP120 and with soluble CD4.^12^ They observed that CD4 bound GP120 did not activate G-protein signaling pathway, acting as an antagonist or inverse against like MRV. In fact, they observed that CD4-GP120 bound CCR5 structure and MRV bound CCR5 structure display few differences on extracellular sites but intracellular signaling protein binding site were similar in inactive conformation. However, there are also various studies showing that GP120 to CCR5 could activate the signaling process^13–15^ or at least bind equally well to G protein coupled or uncoupled CCR5 subpopulations.^16^ These observations lead to the indication that CCR5 displays conformational heterogeneity on cell surfaces and different subpopulations of the receptor could be employed by binding partners.^13, 16–19^ HIV-1 in fact shown to exploit conformational heterogeneity and escapes inhibition by chemokines.^16^ CCR5 conformational heterogeneity is generally associated to its coupling state to G proteins, as expected since CCR5 cycles between active and inactive conformations in this case. The different conformations of CCR5 could also be due to different phenomena such as its oligomerization state^20^, its post-translational modifications^21^ or even the composition of membrane bilayer.^22^ For some GPCRs, the protonation states of ionizable residues displayed to affect the conformational states as shown by several studies.^23–27^ However, to our knowledge the effects of ionizable critical residues of CCR5 was not considered in detail for its conformational heterogeneity.

Molecular dynamics simulations (MD) offer an excellent tool to analyze and sample the conformational states of proteins could undergo. Not only MD provide dynamic structures but also allows to identification of protonation states for ionizable residues. In number of studies, MD simulations have been conducted for CCR5 generally initiated from resolved structures of CCR5 with various ligands. ^28–32^ While the protonation states of all ionizable residues were not mentioned in detail (unless 3D structure was provided as supplementary file), they could be deduced for binding site residues. An important ionizable binding site residue was GLU283^7.39^ (subscript denotes the Ballesteros-Weinstein numbering), as its mutation lead to decreased activity and binding.^28, 33^ In their study, Jacquemard et al.^34^ considered GLU283^7.39^ of CCR5 to be ionized in their MD simulations of CCR5 in complex with various HIV-1 GP120. Salmas et al. ^29^ performed MD simulations of CCR5 in complex with MRV with protonated GLU283^7.39^. In many studies for which a novel inhibitor was searched for CCR5, the protonation state of GLU283^7.39^ was considered as ionized to mimic MRV.^35–36^ On the other hand, Isaikina et al.^28^ performed replica MD simulations with protonated GLU283^7.39^ of CCR5 after determining an active conformational structure of CCR5 in complex with super-agonist [6P4]CCL5. Our investigation of publicly available structures showed that although GLU283^7.39^ was a buried residue, water molecules could still access this region and protonate this residue. In fact there were water molecules resolved in binding site in crystal structure of CCR5 with MRV.^37^ Also, our protonation state prediction of protein site chains assessed GLU283^7.39^ to be protonated when it was in complex with HIV-1 GP120 so that hydrogen bond network could be optimized. Previous studies on GPCRs indicated the effects of pH and ionization state of certain residues on active/inactive state of the receptor as already mentioned in previous paragraph.^23–27^ Hence, it is intriguing to study the possible effect of the protonation state of a binding site residue GLU283^7.39^, whether it could lead to conformational differences especially when CCR5 is in complex with different binding partners. The protonation state of this residue could also have impacts on the design of novel CCR5 inhibitor as it is illustrated that different ligands could lead to titratable residues to be in different protonation states.^38–40^ Hence, consideration of protonation state of this binding site residue could be impactful for drug development and discovery process.

In this study, our aim was to investigate the role of CCR5 binding site residue GLU283^7.39^ in terms interactions with binding partners, as well as its effect on conformational subpopulation of CCR5. For this reason, we have performed long and repeated atomistic MD simulations of CCR5 in complex with HIV-1 GP120 or in complex with MRV or in apo form. The protonation state of CCR5 GLU283 was considered both as ionized and protonated while all other ionizable residues were adjusted to be same for the systems. Our results indicated that there were distinctions in binding modes of GP120 and MRV in case of protonation state changes. CCR5 itself as a receptor with conformational heterogeneity displayed distinct changes in certain motifs of structure especially when it is in complex with GP120.

## 2. METHODS

### Protein Complex Preparation

The recently resolved cryo-EM structures of HIV-1 GP120 in complex with human CCR5 and soluble four domain CD4 receptor was retrieved from protein data bank (PDB IDs: 6MEO at 3.9 resolution).^12^ The structure preparation steps for all protein systems were performed using Maestro program.^41^ As soluble CD4 was used during determination of these cryo-EM structures, the transmembrane and cytoplasmic parts of primary receptor CD4 were missing. Additionally, the extracellular domain of CD4 receptor which has four immunoglobin domains D1-D4 were only available at lower resolution structure (4.5 Å resolution, PDB ID: 6MET). Hence, though we have used the cryo-EM structure refined by Shaik *et al*.^12^ with the higher resolution (3.9 Å resolution, PDB ID: 6MEO), we have taken domains D3-D4 from the structure with lower resolution. The missing transmembrane and cytoplasmic domains (residue numbers 396-458) were previously separately determined (PDB ID: 2KLU)^42^ and these part are added to CD4 extracellular domains in complex with HVI-1 GP120. There were still nine residues (residue numbers 387-396) of CD4 receptor that were not determined by experimental methods, hence this region was modeled using ‘crosslink protein’ options of Maestro program. Additionally, some regions of GP120 envelope protein were unresolved in the cryo-EM structure (PDB ID: 6MEO) such as loop regions between residues 131-185 and residues 396-404 (UniProtKB: Q70145). While the shorter unresolved loop region was modeled using the *de novo* loop modeling of Prime module, structure prediction of the longer loop region was not feasible to be modeled, hence the residues were simply linked together as they were also in close proximity to each other. The structure of the prepared system could be found in Fig. S1. After filling the gap regions of proteins in complex, the protonation states of amino acid residues were again determined using PROPKA^43^ at physiological pH of 7.4. Here we have considered two different protonation states for CCR5 residue GLU283^7.39^, protonated or ionized though PROPKA predicted GLU283^7.39^ to be in protonated state when in complex with GP120 (see Table S2 for predicted p*K*_a_ of this residue). In the last step of protein preparation, a restrained minimization was again applied with OPLS3 force field^44^ to dispose of steric clashes.

The crystal structure of CCR5 in complex with MRV was again taken from protein data bank (PDB ID: 4MBS)^45^. Firstly, the rubredoxin added to third intracellular region (ICL3) as well as solvent molecules away from the vicinity of ligand MRV (> 5 Å) were removed. There were missing residues of ICL3, namely CYS224, ARG225, Asn226 and GLU227 which were again modeled by ‘crosslink proteins’ module of Maestro program.^46^ The mutated residues for thermal stability of the structure were converted to wild type residues based on UniProt sequence (UniProtKB: 51681). The structure of CCR5 coreceptor in MRV bound form only contained residues 19-313, missing the crucial N-terminus (Nt) region for HIV binding. While residues 7-27 of Nt region is solved by NMR and deposited as another structure (PDB ID: 2L87)^47^, residues 1-7 could not be determined due to the flexibility. Nt region residues 7-19 was added to the previously prepared structure after alignment of overlapping regions of two structures while remaining Nt residues 1-7 were modeled in Maestro and again combined with the previous structure. The Nt region which formed an extended loop were minimized by utilizing loop refinement of Prime module.^48^ The newly prepared structure contained 1-313 residues of CCR5 in complex with MRV. After the preparation of CCR5 structure with intact Nt region, the protonation states of residues need to be adjusted. Here, the protonation states of protein side chains at physiological pH of 7.4 were determined using PROPKA.^43^ Two systems with ionized and protonated binding site residue GLU283^7.39^ was prepared. Additionally, it is required to determine the ionization and tautomeric state of the co-crystallized ligand in complex with the protein and here, Epik^49^ is used for this purpose. In the case of ionized GLU283^7.39^ system, the ligand molecule was predicted to be positively charged with tropane group nitrogen atom bearing the charge while this nitrogen atom is considered as neutral in the case that GLU283^7.39^ was protonated (Table S2). The final CCR5-MRV complexes structure with predicted ionization states were then optimized by restrained minimization on heavy atoms with OPLS3 forcefield.^44^ We have also performed MD simulations of CCR5-MRV complex in which the starting coordinates of protein structure taken from CD4/GP120 bound CCR5 (PDB ID, 6MEO).

The recently released *apo* form structure of CCR5 in complex with G_i_-protein (PDB ID:7F1S) was downloaded and prepared using the following steps. The structure was in active form as it was in complex with coupling protein, but we were interested in the conformations CCR5 by itself can undergo. Hence, G_i_ protein was deleted and missing loop regions such as between residues 91-95 were modeled using Prime module of Schrodinger. The Nt region, residues 1-32, was also not present, other CCR5 structures were used: residues 17-32 were taken from another CCR5 structure (PDB ID, 7F1Q) and remaining 1-16 were taken from already prepared *holo* form structure. At the end, an *apo* form CCR5 structure with residues 1-313 was obtained. We also initiated MD simulations in which the apo form of CCR5 is obtained by removing ligand MRV or CD4-GP120. PROPKA^43^ was used to determine the protonation states of residues for *apo* CCR5 receptor. Again, two different ionization states were considered for residue GLU283^7.39^. As a final preparation step, restraint energy minimizations and geometry optimizations with OPLS3 forcefield^44^ were applied. Table S2 displays the information about the prepared systems along with pKa value of GLU283^7.39^ in different systems predicted by PROPKA.

### System Preparation and Molecular Dynamics Simulations

Both CCR5 and CD4 receptors were embedded into 1-Palmitoyl-2-oleoyl-sn-glycero-3-phosphoethanolamine (POPE) membrane bilayer. The orientations of proteins in the lipid bilayer were adjusted using Orientations of Proteins in Membranes (OPM) server ^50^ and special care was taken to embed only the transmembrane part of CD4 receptor. The membrane embedded protein systems were placed in orthorhombic boxes of TIP3P^51^ water molecules with box sizes calculated based on buffer distance which were chosen as 10.0 Å along all three dimensions. The systems were all neutralized and ionic concentration of 0.15 M NaCl solution was used to adjust concentration of the solvent systems. Desmond program^52^ was utilized for all atom MD simulations. Before starting the production MD simulations, relaxation protocols were performed to equilibrate the systems and membrane relaxation protocol of Desmond as suggested was applied. During production simulations, the initial temperature was set as 310 K and controlled using Nose-Hoover thermostat.^53–54^ The pressure was set as 1.01325 bar and it was controlled by the Martyna-Tobias-Klein method^55^ with anisotropic pressure coupling. To calculate the equation of motions in dynamics, RESPA a multi-step integrator with time steps of 2.0, 2.0 and 6.0 fs for bonded, *near* non-bonded and *far* non-bonded interactions respectively was utilized. The cut-off value for the short-range electrostatic and Lennard–Jones interactions was set as 9.0 Å. Particle mesh Ewald (PME)^56^ method along with periodic boundary conditions (PBC) was used to estimate the long-range interactions. All systems were subjected to 500 ns MD simulations with three independent replicate simulations and every 100 ps intervals of MD simulations were written as trajectory file for analysis.

### Analysis of Molecular Dynamics Simulations

#### Root Mean Square Deviations (RMSD) and Root Mean Square Fluctuations (RMSF)

The trajectory files collecting the positions of each atom in production MD simulations were used for analysis of atomic deviations observed in a time-dependent manner. Root mean square deviations (RMSD) and root mean square fluctuations (RMSF) analysis were performed to assess the stability of protein structures. For each replica MD simulation, RMSD and RMSF analysis were performed separately and each frame in trajectory files were first aligned to the structure obtained after protein preparation step, i.e. initial structure, explained in the previous section. RMSD values were calculated by considering each protein as separate chains and after alignment to only the initial chain structure. For CD4 receptor, RMSD of domains 1 (DM1) and domain 2 (DM2) were also calculated separately for MD simulation time. For the calculation of RMSF values, the proteins in CD4-CCR5-GP120 complex are considered as separate chains and for each chain, the frames in MD trajectories were aligned to initial chain structures. In both RMSD and RMSF analysis, carbon alpha (Cα) atoms of proteins were used for alignment and atomic deviations or fluctuation calculations.

#### Contact Analysis

The non-covalent interactions between GP120 glycoprotein and CCR5 coreceptor were analyzed using GetContacts application, a python-based code obtained from (https://getcontacts.github.io/). MD trajectories from each independent replica simulations of each system (Simulation IDs: MD-1 and MD-2) were concatenated and the frequency of intermolecular interactions were computed in trajectory-based approach. In the study, salt-bridge interactions, hydrogen bonds and π-cation interactions were considered and quantified separately. The details about the geometric and chemical criteria for each interaction type could be found in the webpage of the code (https://getcontacts.github.io/interactions.html). The calculated interaction frequencies were converted to percent values in which 60% value of frequency represent that specific interaction was maintained during 60% of the simulation time.

#### Cluster Analysis

The concatenated MD trajectories for each system were used in cluster analysis and Trusty Trajectory Clustering (TTClust)^57^ python program was employed for this reason. The centroid clustering method was used, and 10 clusters were requested. The protein structures were aligned based on Cα atoms. RMSD and distance matrix were also calculated for carbon alpha atoms. Representative structures for each cluster were also generated to be used in visualizations. We performed clustering for GP120-CCR5 complex as well as MRV bound CCR5 in addition to clustering for only CCR5 chains in the systems.

#### Principle Component Analysis (PCA) and Cross Correlation

We used Bio3d package^58–59^ which is a platform independent R package. The trajectories obtained from independent MD simulations were concatenated and frames were aligned with respect to initial (reference) frame. Only Cα of CCR5 chains in each system were considered as the aim was to compare the conformational differences in CCR5 in different systems. The detailed explanations of the analysis were given in our previous papers^60–61^ as well as the package manual (http://thegrantlab.org/bio3d_v2/user-guide). Briefly, for PCA, the covariance matrix, *COV_i,j_* was generated by considering cartesian coordinates of each Cα atom and their displacements from average positions. The diagonalization of this matrix provided eigenvectors, referred as principal components (PC) and their corresponding eigenvalues which gives the variance of the distributions along PCs. By PCA, we could identify intrinsic motions in protein structures which were usually canceled by random motions.^62^ For cross correlation analysis, the normalized covariance matrix of atomic fluctuations, *C_i,j_* was calculated and a graphical representation of cross correlation coefficient (obtained from covariance matrix) was generated. As the matrix is normalized, the values of *C_i,j_* varies between −1 to 1 with *C_i,j_* = 1 representing completely correlated motions (same period and same phase), *C_i,j_* = −1 representing completely anticorrelated motions (same period and opposite phase while *C_i,j_* = 0 indicating motions are uncorrelated.^63–64^ We generated dynamic cross correlation maps (DCCM) to visualize this matrix. By cross correlation analysis, we can assess the extent of correlations in atomic displacement and determined dynamical effects of binding partners on these displacements. In practice, to obtain PCs and eigenvalues we used pca.xyz() command to while using dccm.xyz() command to obtain DDCMs.

#### Distance, Torsional and RMSD Analysis

Bio3D package^58–59^ was again utilized to compute the distances between specific atoms in protein structures. Once again, MD trajectories of independent replica simulations were concatenated. dist.xyz() command was applied along with the atom indexes for analysis. The torsion angles for side chains of specified residues were also calculated using Bio3D package^58–59^, torsion.xyz() command was used along with the atom indexes for which torsional analysis performed. We also calculated the RMSD values of important motifs with respect to inactive structure by utilizing Bio3D package with rmsd() command along with indexes of Cα atoms in those motifs.

## 3. RESULTS AND DISCUSSION

### MD Simulations of CCR5 in Complex with Different Binding Partners

MD simulations were repeated three times for all the studied systems. As can be seen from Table S1, we have 10 systems in which CCR5 was either in complex with GP120, MRV or in *apo* form and binding site residue, GLU283^7.39^ was either protonated (GLH283) or ionized (GLU283). In total, we performed 15.0 µs MD simulations and collected the simulation trajectory frames for analysis. We first evaluated RMSD of protein carbon alpha (Cα) atoms displayed for each system. For CD4-GP120 bound CCR5 systems, RMSD values were calculated for the whole protein complex as well as for separate protein chains (Fig. S2). We have observed different behavior for independent MD simulations though all protein structures displayed stability in certain time courses of MD and the changes observed were gradual in general. The protein complex CD4-GP120-CCR5 complex displayed very large RMSD values (larger than 10.0 Å), hence high fluctuations were observed compared to initial structure during MD simulations. Our analysis indicated that this was due to the change in the orientation of CD4 receptor with respect to membrane mostly. CCR5 protein by itself displays also considerably smaller fluctuations compared to whole complex (around 6.0 Å RMSD values), and it is the loop regions, specifically N-terminal loop region that had larger changes as can be seen low in RMSF plots (Fig. S3, CCR5 plots). GP120 protein by itself displays also relatively smaller RMSD values, and in this case, it was again variable loops (V1-V3) that have higher fluctuations (see Fig. S3, GP120 plots). As mentioned, CD4 receptor was undergoing orientational change with respect to membrane bilayer. Hence, the RMSD values for CD4 receptors were substantially large (over 10.0 Å), and a sharp increase was observed especially during the early stages of simulations. This behavior is expected as we have modeled the missing parts of CD4, and it required same time for structure to equilibrate. When we have analyzed and compared the trajectory frames specifically for CD4 receptor structural changes, we have seen that the extracellular domains D3-D4 display high RMSD compared to initial conformation due to their movement towards to the membrane during simulations (Fig. S2, CD4 D1 & D2 plots and S3, CD4 plot). Nevertheless, RMSD plots for replica simulations with both protonated and ionized GLU283 reached a plateau after the initial sharp increase for CD4 receptor.

The RMSD values of CCR5 receptor in complex with MRV displayed moderate RMSD values for MD simulations (around 4.0-6.0 Å) with values being higher for the system with ionized GLU283 (Fig. S4). Though we have seen again it was loop regions with higher fluctuations as TM regions have values less than 2.0 Å (Fig. S5). Similar behavior observed of MD simulations of *apo* systems which were initiated from different 3D structures (from PDB entries; 7F1S, 6MEO and 4MBS). For *apo* simulations initiated from active CCR5 structure (PDB entry 7F1S), we observed quite large initial RMSD values (Fig. S6) which were due to N-terminus regions as it was modeled from other structures (see Fig. S7, RMSF plots). The same observation was true for the *apo* MD simulations initiated by removing CD4 and GP120 proteins from CD4-GP120-CCR5 complex (PDB entry 6MEO), but this time it was due to the loss of interactions with GP120 that caused N-terminus of CCR5 to fluctuate (Fig. S8 and S9). The *apo* CCR5 simulations initiated by removing co-crystalized MRV compound (PDB entry 4MBS) displayed lower RMSD values compared to other *apo* simulations though still N-terminus region had the highest fluctuations (Fig. S10 and S11). All in all, the systems simulated had explainable RMSD values and gradual changes were observed after the initial MD times.

### Interactions Between GP120 and CCR5 for Ionized and Protonated Forms of GLU283

As mentioned before, the V3 loop of GP120 glycoprotein specifically interacts with the residues in chemokine recognition site 2, CRS2 of CCR5.^12^ Additionally, there are important interactions formed between GP120 and N-terminus of CCR5, especially sulfated tyrosine residues, TYR10 and TYR14, are critical for recognition of the virus. The interaction network between GP120 and CCR5 observed in cryo-EM structure was shown in the paper of Shaik *et al*.^12^ However, these interactions could only be observed in a static manner by crystal or cryo-EM structures. We computed the frequencies of non-covalent interactions such as salt-bridges, hydrogen bonds and π-cation interactions between GP120 and CCR5 for the systems with different protonation state of binding site residue GLU283^7.39^ (Fig. S12). There were observed differences in frequencies of protein-protein interactions for the systems. These differences were as expected most pronounced for CCR5 GLU283^7.39^ residue and its network of interactions with GP120. We displayed in detail the critical interactions between these two proteins that were also specified by Shaik *et al*.^12^ with ionized and protonated GLU283^7.39^ of CCR5, respectively (Fig. S13 and S14, respectively).

We initially checked the interactions between V3 loop of GP120 and CCR5. The PRO311 of V3 loop that was indicated to be reaching into CRS2 of CCR5 and found between side chains of Trp86^2.60^ and TYR108^3.32^ in cryo-EM structure. This residue of V3 loop was well packed in this region throughout MD simulations. On the other hand, we observed that sidechain of ARG313 of V3 loop which appeared between TYR25^6.51^ and GLU283^7.39^ of CCR5 in cryo-EM structure has moved away from this area specifically with GLH283^7.39^ of CCR5. In this case, ARG313 of V3 loop formed persistent (i.e. interaction frequency higher than 60%) salt-bridge and hydrogen bond interactions with ASP276^7.31^ of CCR5. This V3 loop residue also formed considerably persistent hydrogen bond with ASN258^6.58^ of CCR5. Related to the side chain re-orientation of V3 residue ARG313, GLY312 and ALA314 were also interacted differently with CCR5 coreceptor in systems with different protonation states of GLU283^7.39^. While V3 loop GLY312 maintained hydrogen bond interactions with GLU283^7.39^, this hydrogen bond interaction was absent with GLH283^7.39^. On the other hand, V3 loop ALA314 formed hydrogen bond with GLN280^7.35^ for at least half of the simulation time for GLH283^7.39^ system, but this interaction was observed scarcely when GLU283^7.39^ was ionized. The interaction networks for V3 loop residues HIS308 and TYR316 with CCR5 residues TYR89^2.63^ and ECL2 residue CYS178 were also slightly different for the two systems in consideration. In system with GLH283^7.39^, V3 loop residue HIS308 hydrogen bonded with CYS178 while V3 loop residue TYR316 hydrogen bonded with TYR89^2.63^. In system with GLU283^7.39^, both HIS308 and TYR316 forms hydrogen bonds TYR89^2.63^ of CCR5 with almost no contact between HIS308 and CYS178. We can also see that some non-covalent interactions were persisting interactions for both systems. Specifically, salt-bridge between extracellular 2 (ECL2) residue GLU172 of CCR5 and ARG304 of GP120 was a well-maintained interaction throughout MD simulations. Another maintained hydrogen bond interaction was between ECL2 residue of CCR5 SER180 and ILE307 of GP120. The hydrogen bond between ECL2 residue SER179 of CCR5 and SER306 of V3 loop was similarly maintained in both systems with different protonation states of GLU283^7.39^ though the interactions were not maintained to large extent.

The interactions GP120 forms with CCR5 Nt critical residues TYR10, TYR14 and TYR15 were mostly similar for differently protonated systems with little variations in interaction frequencies. Both sulfonated tyrosine residues did not form persistent salt-bridges; TYR10 only formed transient salt-bridge with LYS416 of GP120 in both systems while TYR14 formed short-lived salt-bridges with ARG298 and LYS435 of GP120. However, the hydrogen bonds sulfonated TYR10 formed with mentioned residues of GP120 were more persistent (frequencies minimum 44%). Additionally, TYR10 of CCR5 formed hydrogen bonds with GLN417 and ILE418. This sulfonated residue also forms π-π interactions with ARG326 of GP120 for both systems as mentioned by Shaik et al.^12^ though not very persistently which could be because it also forms π-π interactions with another close by residue ARG414 of GP120. TYR14 of CCR5 also formed hydrogen bond interactions with backbone atoms of GLY436 in both GP120 bound system. This sulfonated TYR14 also involved in π-π interaction with LYS435, and there was substantial a difference in frequencies (larger than 20%) for differently protonated systems as this interaction was more maintained in protonated GLU283 CCR5 system. TYR15, which was not sulfonated, formed permanent hydrogen bonds with backbone O atom of PRO433 in GP120. It also involved in π-π interaction with LYS207 of GP120 though the interaction was not persistence especially in system with ionized GLU283 CCR5. Other Nt residues of CCR5, LYS22 and LYS26, were also forming contacts with GP120, specifically with ASP320 residue. In the system containing ionized GLU283 CCR5, it was LYS22 that was mainly forming hydrogen bonds and salt-bridges with ASP320. But in the system containing protonated GLU283 CCR5, both LYS22 and LYS26 were involved in forming both hydrogen bonds and π-π interaction. In total, the interactions formed between GP120 and CCR5 were more maintained interactions when CCR5 contained protonated GLH283^7.39^.

The binding modes of V3 loop for ionized and protonated GLU283^7.39^ systems were also compared (Fig. 1). Representative structures obtained after cluster analysis were used to represent differences in binding modes. The structures were superimposed on top of each other by aligning the CCR5 chain. We observed that V3 loop was flexible in the binding cavity hence the orientation of it was not identical for two systems. Though PRO311 of GP120 was oriented the same, it was ARG313 side chain that was oriented differently in the binding site of two systems (Fig. 1b and 1c). We also observed differences in flexible extracellular loops ECL2 and ECL2 for two systems which could indicate that V3 differently shapes these sites of CCR5 coreceptor. These comparisons indicate that the ionized and protonated form of GLU283^7.39^ in CCR5 lead to different binding modes for V3 loop of GP120. Also, the extracellular site of CCR5 itself is differently shaped which could affect the allosteric network within the receptor and could even lead the receptor to sample an active-like structure (discussed in later sections).

**Figure 1.**
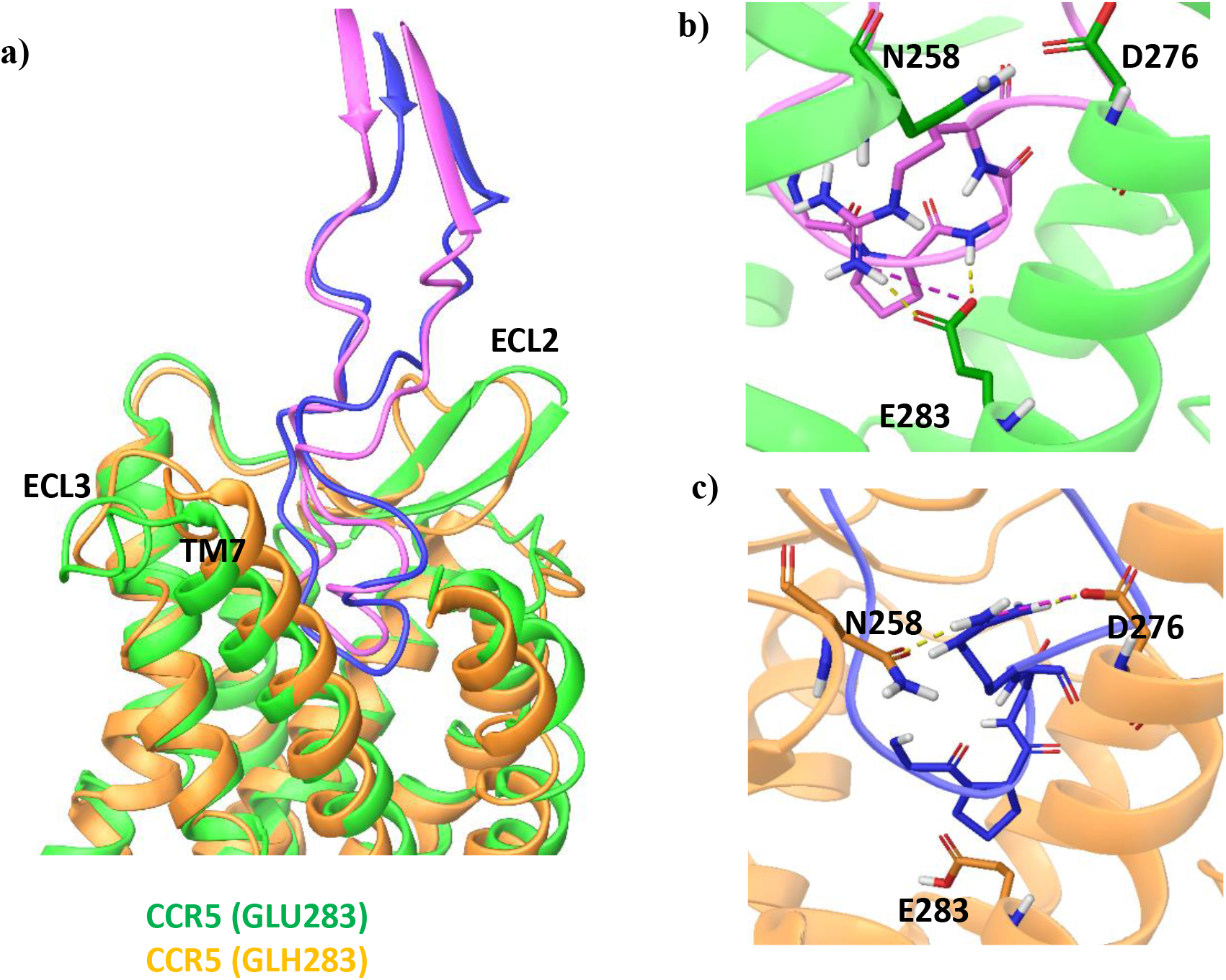
CCR5 structures in complex with CD4-GP120 in systems with protonated and ionized GLU283. Only V3 loop of GP120 is displayed and Nt of CCR5 is omitted for better clarity. Representative structures from clusters are used. a) Comparison of V3 loop binding modes in differently protonated CCR5 systems. b) Detailed interaction networks that were significantly varied in the systems. Yellow dashed lines represent hydrogen bonds, pink ones represent salt-bridge

### Interactions Between MRV and CCR5 for Ionized and Protonated Forms of GLU283

MRV is the only approved CCR5 inhibitor as mentioned before. This molecule was suggested to be an allosteric inhibitor^37^ though Shaik et al. argued that it is a competitive inhibitor as the binding site of MRV overlaps partially with binding site of GP120. Additionally, MRV was characterized as an inverse agonist that retain CCR5 in inactive conformation.^65^ Tan et al.^37^ resolved the structure of CCR5 in complex with MRV and verified that the receptor was in an inactive state. Our MD simulations initiated from this inactive state of CCR5-MRV complex, and we did not observe high changes in the TM regions of the receptor throughout MD simulations (Fig. S4 and S5). Hence, we can say that the receptor maintained its inactive conformation for both systems with ionized and protonated GLU283^7.39^ CCR5.

The interaction profile of MRV with differently protonated CCR5 was not same based on the dynamical interaction analysis evaluated from concatenated MD trajectories (Fig. S15 and S16). It should be noted that there was not only the difference in protonation state of CCR5 residue GLU283 for the systems but also the ionization states of MRV were different; nitrogen atom of tropane was either positively charged (for system in GLU283 was ionized) or neutral (for system in GLU283 was protonated). Hence, the total charges for both systems of differently protonated CCR5 were same. These systems with different protonation and ionization states were compared in terms of the protein-ligand interactions. The MRV-CCR5 interactions were again deduced from concatenated MD trajectories though in this case simulation interaction analysis of Maestro was used to compute the persistency of interactions. MRV maintained most of the interactions observed in crystal structure^37^ during MD simulations. Specifically, following CCR5 residues maintained hydrophobic contacts: TRP86^2.60^, TYR89^2.63^, TYR108^3.32^, PHE109^3.33^, PHE112^3.36^, ILE118^3.42^, TRP248^6.48^ and TYR251^6.51^ though to frequencies of interactions varied for the systems (Fig. S15a and S16a). Trp86 engaged in a π-π interaction with MRV for the system with ionized GLU283 of CCR5 while this interaction was absent in GLH283 system as MRV does not have a positively charged nitrogen atom there (Fig. S15b, c and S16b, c). TYR108^3.32^ formed both π-π and π-cation interactions with MRV in system with ionized GLU283 but it only formed π-π interaction in system with protonated GLU283. The hydrogen bond interaction with TYR37^1.39^ was well-maintained (over 60%) though hydrogen bond formed with TYR251^6.51^, that was present in crystal structure, was short-lived for both systems (Fig. S15b, c and S16b, c). MRV engaged in hydrogen bond interactions with GLU283^7.39^ of CCR5 for both systems with similar interaction frequencies though the atoms involved in the interactions differed. MRV formed more maintained interactions with CCR5 when GLU283^7.39^ was ionized.

We also compared the binding modes of MRV in CCR5 binding pocket with ionized and protonated form of GLU283^7.39^ (Fig. 2). We again considered representative structures from clusters to indicate differences in structures and MRV binding modes. MRV was also flexible in CRS2 binding site of CCR5 and oriented itself slightly differently in the compared systems. Specifically, the phenyl ring that was inserted deep into the cavity was not similarly positioned. There were also variations in the conformation of certain regions of CCR5, especially ECL3 loop. There were also differences in conformation of TM6 and TM7 extracellular sites as they could not be aligned though other TMs did. These differences between systems indicated that the binding mode of MRV is affected by the protonation state of active site residue GLU283^7.39^ as expected. It also showed that initiating MD simulations with different protonation states for this residue could lead to changes in conformations of CCR5 sampled.

**Figure 2.**
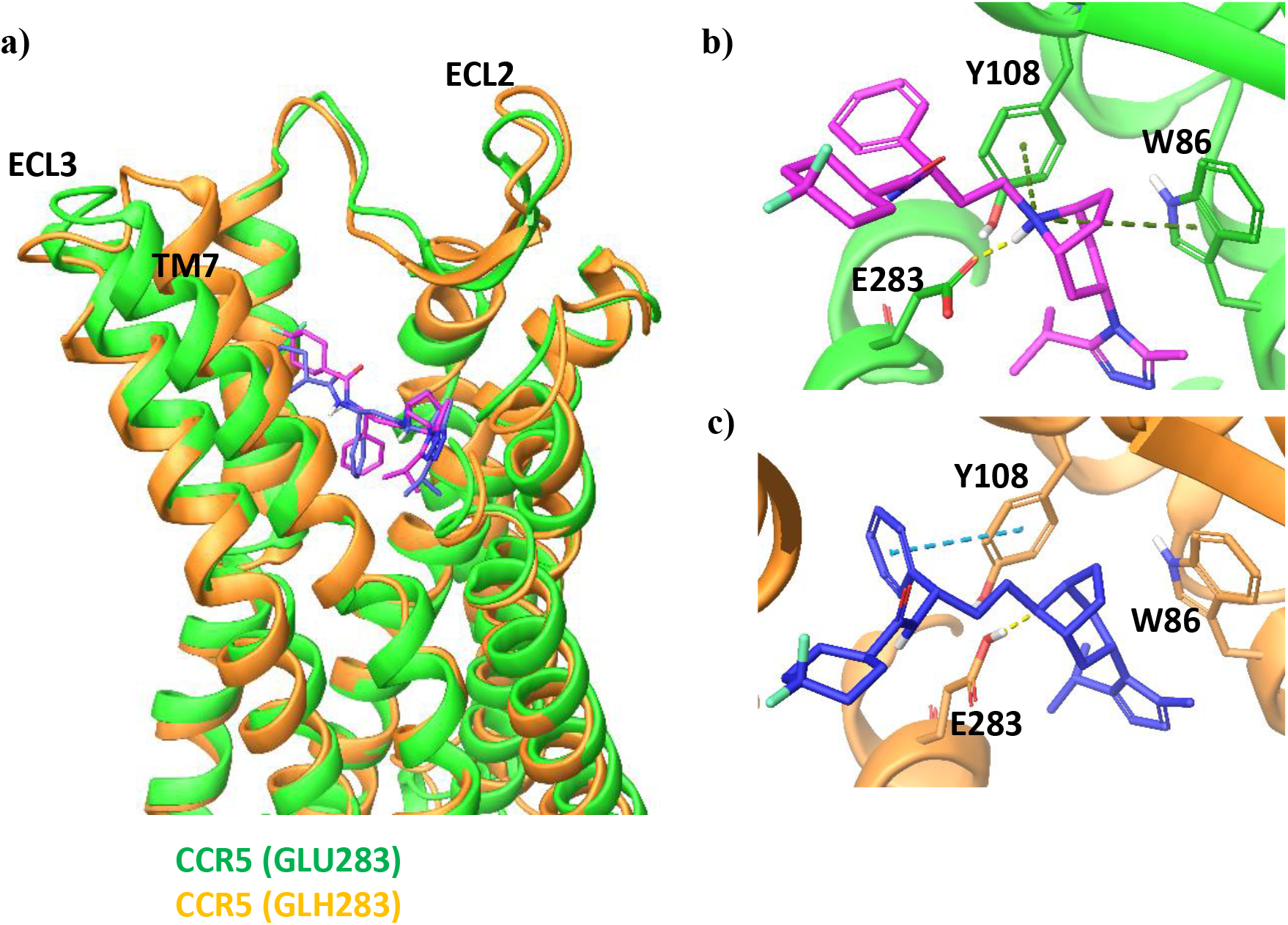
CCR5 structures in complex with MRV in systems with protonated and ionized GLU283. Nt of CCR5 is omitted for better clarity. Representative structures from clusters are used. a) Comparison of V3 loop binding modes in differently protonated CCR5 systems. b) Detailed interaction networks that were significantly varied in the systems. Yellow dashed lines represent hydrogen bonds, pink ones represent salt-bridges. Yellow dashed lines represent hydrogen bonds, blue ones represent π-π interactions, green ones π-cation interactions.

### Differences in Binding Modes to CCR5 for GP120 and MRV

As mentioned, the interactions between GP120 and CCR5 were more maintained in system with GLH283^7.39^ of CCR5 while for MRV it was the system with GLU283^7.39^ that leads to formation of more maintained interactions of it with CCR5. The predicted p*K*_a_ values of CCR5 GLU283^7.39^ was also indicating the same (Table S2). Hence, we decided to compare the binding modes of GP120 and MRV with the systems in which they formed more-maintained interactions, meaning that for CCR5-GP120(/CD4), the system with GLH283^7.39^ was considered while for MRV, the system with GLU283^7.39^ was considered. The representative structures for both systems indicated that there were differences in extracellular sites ECL2 and ECL3 as expected but TM regions aligned on top of each other with good agreement (RMSD = 1.72 Å, Fig. 3a). There were also no significant differences in the intracellular sites of TM regions. As mentioned by Shaik et al., the binding sites of GP120 of HIV-1 and MRV overlap especially V3 loop of GP120 occupies partially a region that MRV binds to (Fig. 3b). When we focused on the binding site, we found out that there were changes in the conformations and orientation (Fig. 3c). As expected, GLU(H)283 of CCR5 oriented differently in the compared systems. TRP86^2.60^ and TYR89^2.63^ of two systems also could not be aligned and displayed dissimilar orientations. Other binding sites residues though had only slightly different conformations and their orientations in CCR5 receptors were similar (Fig. 3c).

**Figure 3.**
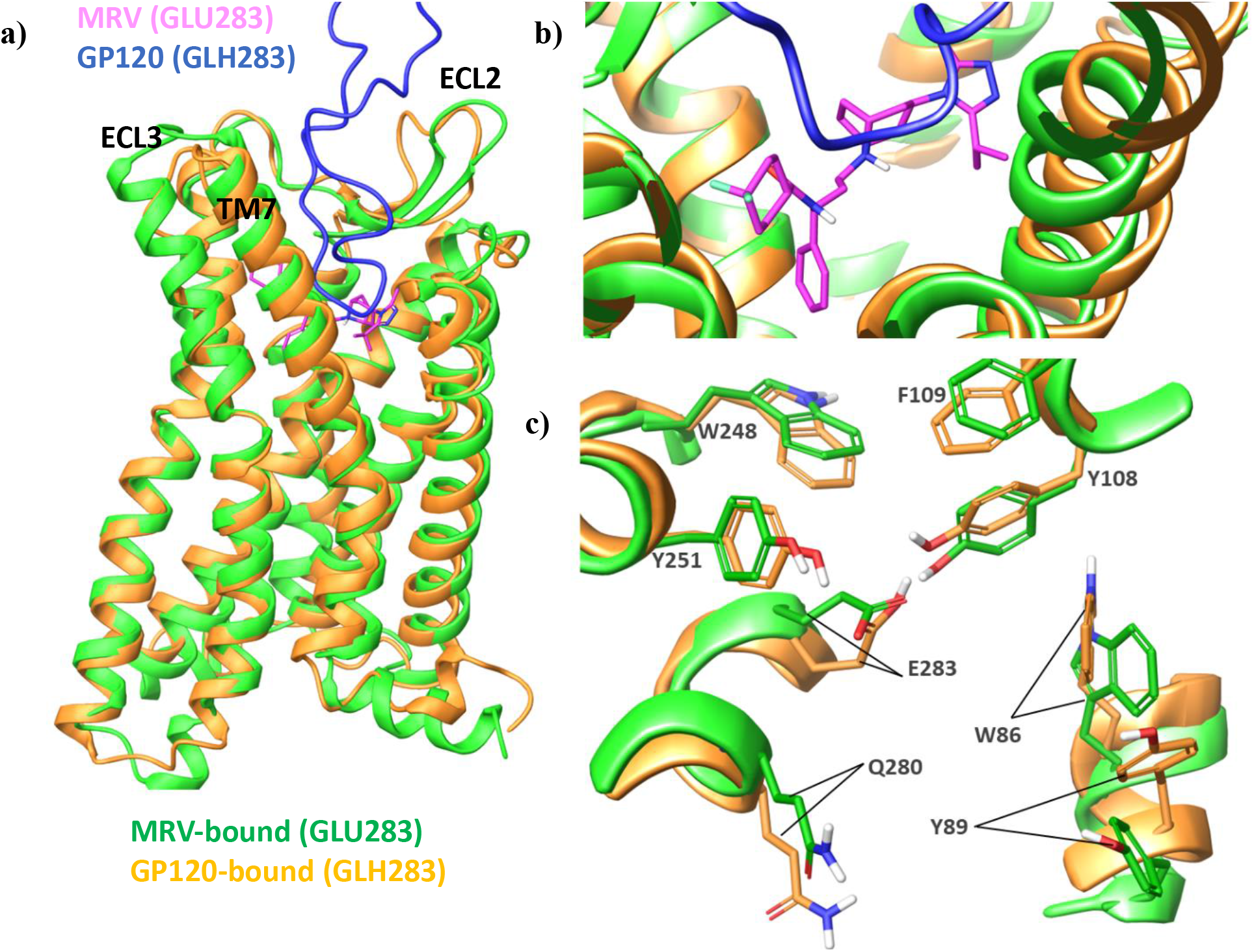
GP120 is displayed and Nt of CCR5 are omitted for better clarity. Representative structures from clusters are used. a) Alignment of structures on top of each other b) Binding site comparison for structures c) Side chains conformational comparison.

### Intra-receptor interactions within CCR5 in Bound and Apo Forms

We analyzed the differences observed in the contacts within the CCR5 receptor while it is bounded to either GP120 or MRV or in *apo* form in different protonation states of GLU283^7.39^. Another objective of ours was to see if there were changes in intra-receptor interactions within CCR5 with the protonation state change of GLU283^7.39^. Hence, for all the systems for which MD simulations were conducted, intra-receptor interactions such as salt-bridges, hydrogen bonds and π-π or π-cation interactions were dynamically evaluated by again GetContacts algorithm (Fig S17). Here, hydrogen bond interactions were calculated for residues that are at least five residues away from each other to avoid considering hydrogen bonds due to helical structure and only displayed the interactions that differ or are indicated to be important in previous studies.

The intra-helical salt-bridge interactions between ASP125^3.49^ and ARG126^3.50^ were differently conserved in the studied systems (Fig S17). This interaction along with the ionic lock of ARG126^3.50^ from with residue at 6 x 30 were stated to broken during activation of the receptor.^66^ In the case of CCR5 there was no ionic lock ARG126^3.50^ forms with residue at 6×30 to stabilize its inactive conformation. We observed the release of ARG126^3.50^ from intra-helical salt-bridge with ASP125^3.49^ for GP120 bound and *apo* states (initiated from PDB 7F1S) as the interaction was maintained less that %50 of simulation time though in systems with GLH283^7.39^ the interaction was comparably more reserved. The salt-bridge interaction between ASP125^3.49^-ARG126^3.50^ was mostly maintained in MRV bound systems which could be helpful in stabilizing these structures in inactive form. There were other the salt-bridge interactions that differ for systems such as ARG230^6.30^ – GLU227^ICL3^ and ARG232^6.32^ – GLU302^8.48^ though it should be noted that for HOLO and *apo* systems (initiated from PDB 4MBS), the ICL3 regions were missing and were modeled hence, the salt-bridge may not be formed.

When we compared the intra-receptor hydrogen bond interactions, the first one we focused on was the hydrogen bond network ASP76^2.50^ residue formed. The protonation state of residue was indicated to be playing an important role in the activation process with the population of protonated residue increasing as the receptor is activated.^2, 23, 67^ In our protein preparation steps, this residue was predicted to be protonated to optimize the hydrogen bond network for all considered systems and hence, we performed our simulations with protonated ASP76^2.50^. Though in CCR5 and chemokines this residue site was not considered for sodium ion binding as resolved structure do not indicate a sodium ion density^2^, still this residue was still conserved in chemokines. In our case, the evaluated dynamics contacts revealed that ASP76^2.50^ forms long lasting hydrogen bonds with two asparagine residues ASN48^1.50^ and ASN293^7.49^ in *holo*, *apo* (initiated from PDB 7F1S, initially active receptor) and GP120 bound case with protonated GLU283^7.39^ (Fig. S17). In the case of GP120 bound to CCR5 with ionized GLU283^7.39^, there was a considerably well-maintained hydrogen bond formed between ASP76^2.50^ and TYR297^7.53^ which was absent in other systems. TYR244^6.44^ was another residue involved in hydrogen network differently in simulated systems. In *apo* state (initiated from PDB 7F1S, initially active receptor), it interacted with GLY202^5.46^ persistently regardless of GLU283^7.39^ protonation state and with ALA249^6.49^ when CCR5 has protonated GLH283^7.39^. But TYR244^6.44^ did not form hydrogen bond with ASN293^7.49^ for *apo* active state systems. On the other hand, bound CCR5 and *apo* state simulations initiated from inactive conformations (from PDB 6MEO or 4MBS) generally had hydrogen bonds between TYR244^6.44^ and ASN293^7.49^. But for these systems, TYR244^6.44^ scarcely formed hydrogen bonds with GLY202^5.46^ or ALA249^6.49^ except for when CCR5 with GLH283 bound to MRV and *apo* simulations initiated from MRV bound structure (Fig. S17).

When we evaluated π-π interactions within CCR5 receptor, there were only a few persistent interactions and they usually involved residue HIS289^7.45^ that is found in the central TM region of the receptor. In GP120 bound CCR5 structures there were persistent π-π bonds formed between HIS289^7.45^ and TYR244^6.44^ but for other systems, this interaction was not well maintained. Other interactions HIS289^7.45^ formed were more transient for all considered systems. There was only one π-cation interaction that in some systems were formed and maintained mediocrely and it was between ECL2 loop residues HIS181 and ARG168. This interaction was also differently maintained for the considered system and Considering that ECL2 region is affected by ligand binding the difference was not surprising.

### CCR5 Conformational Differences in Bound and Apo Forms

Another aim of ours was to expose any changes/differences observed in CCR5 receptor conformation, especially related to receptor activation, when the receptor was in complex with binding partners, GP120 and MRV or in its *apo* state as well as when the protonation state of GLU283^7.39^ differs. As CCR5 is a member of GPCR family that is well studied structurally and dynamically, there are already some distances and motifs that are indicated to be important for receptor structure and activation.^28, 32^ Hence, we utilized the trajectories obtained from MD simulations to determine changes in these distances and motifs as well as considering other important activation switches of class A GPCR indicated in other studies. Here, we focused on and compared the changes in bound and *apo* form of CCR5 and for bound forms we specifically considered the protonation state of GLU283^7.39^ based on the interaction profile between CCR5 and binding partner. In other words, for GP120 bound CCR5, we considered the system in which GLU283^7.39^ was protonated while for MRV bound CCR5, we considered the system in which GLU283^7.39^ was ionized since these systems resulted in more well-maintained interactions as discussed in previous parts. For *apo* systems, we considered the simulations initiated from active CCR5 structure (from PDB 7F1S) and systems with both protonation states of GLU283^7.39^ were considered for comparison.

We initially evaluated whether there was any state change in CCR5 structure related to its activation. Hence, we calculated a distance change, Δ*d*, recommended previously by database GPCRdb and utilized in one of our previous studies ^68^ for another class A GPCR. This Δ*d* distance was calculated by considering Cα distances between certain residues at specific positions and indicates the changes in the intracellular side of TM2-TM6 and TM3-TM7 was calculated dynamically. For CCR5, firstly the Cα distance between ILE67^2.41^ and PHE238^6.38^ was measured and from this distance we subtracted the Cα distance between ILE120^3.44^ and ILE296^7.52^. The receptor structures classification was suggested to be as follows: inactive conformations if Δ*d* was smaller than 2.00 Å, active conformation if Δ*d* was larger than 7.15 Å and an intermediate conformation if the value is between these two limits. The distribution of the distance was also plotted for different systems for comparison (Fig. 4a and Fig S18). Based on this analysis, we can say that MRV bound CCR5 structures mostly adopted inactive conformations for both protonated and ionized GLU283^7.39^ as the Δ*d* values did not deviate from 2.00 Å for all MD trajectories. However, we observed a different case for GP120 bound CCR5 structures for differently protonated GLU283^7.39^ systems. Specifically, in protonated GLU283^7.39^ system, GP120 binding could cause conformational change in CCR5 such that it can start adopting an active-like state. This behavior was not observed for GP120 bound CCR5 with ionized GLU283^7.39^ or in *apo* systems for which simulations were initiated from bound state conformations (PDB IDs 6MEO and 4MBS, see Fig. S18). In fact, comparison of structures taken from MD simulations also show that GP120 bound CCR5 structure can adopt a conformation on the intracellular TM region that was comparable to active state one, specifically (Fig. 4b). Hence, we can see the effect of protonation state for GLU283^7.39^ on CCR5 conformational state for GP120 binding.

**Figure 4.**
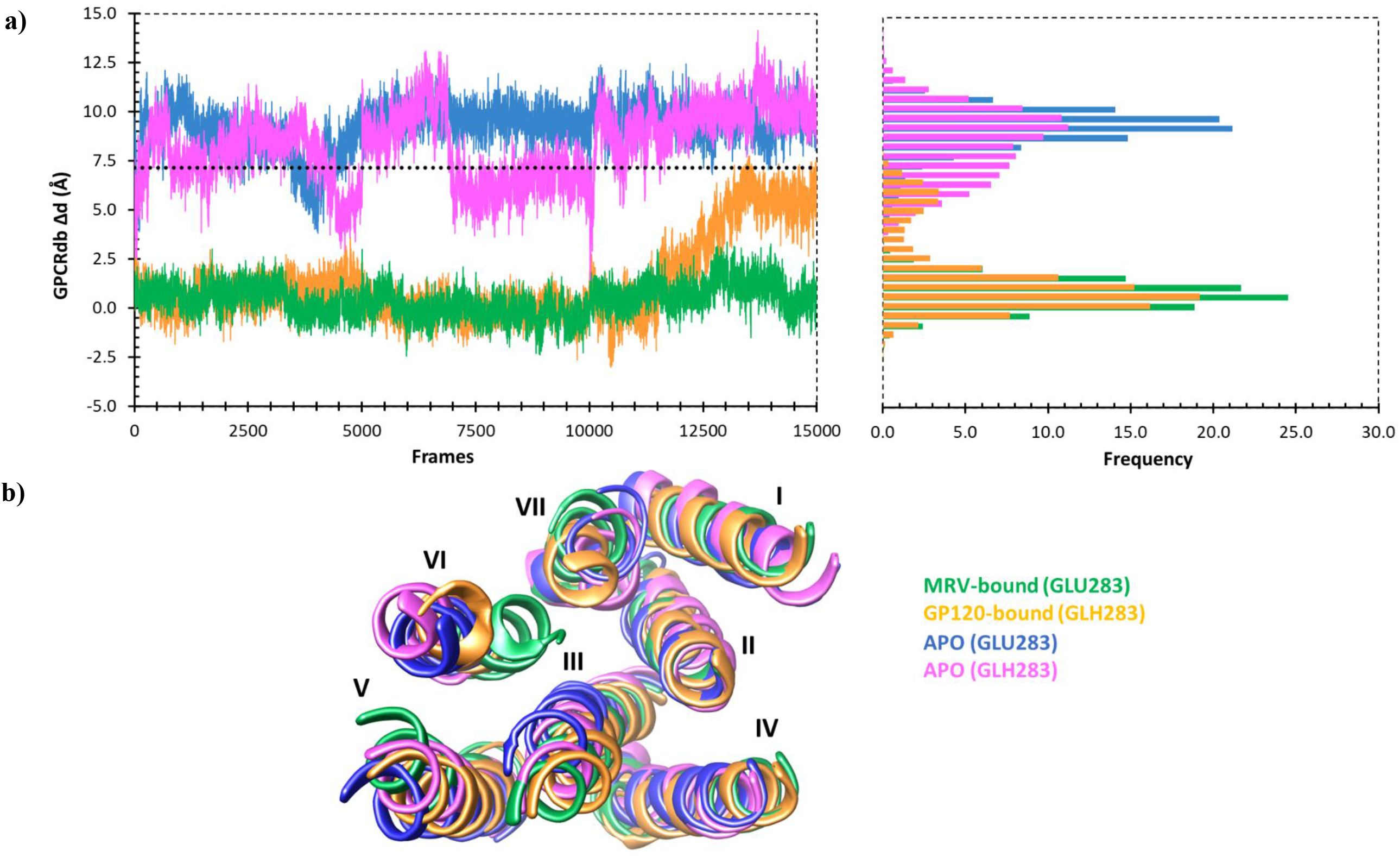
Classification of CCR5 receptor state as active, intermediate and inactive. a) Δ*d* distance measured for the systems using concatenated MD trajectories and frequency distribution of the distance. b) comparison of TM conformations on the intracellular sites. The representative structures of largest clusters were taken for *apo* state while for MRV and GP120 bound states, CCR5 structures with largest distance from MD frames were chosen.

We also considered other distance changes associated with class A GPCR receptor activation. One commonly considered distance was between TM3 and TM6, specifically corresponding to distance between ARG126^3.50^ – VAL234^6.34^ in CCR5.^69^ The values of distance over 9.70 Å was assumed to indicate GPCR adopting active state. Our evaluation of the Cα distance between these two residues (Fig. S19) lead to a similar outcome as Δ*d* distance evaluation. Again, GP120 bound CCR5 with GLH283^7.39^ can adopt an active-like conformation (at least in some of MD frames) with TM3-TM6 distances reaching over the cut-off value of 9.70 Å. In this aspect, this system resembled simulations started from *apo* active state. The CCR5 conformation in other systems, especially MRV bound ones, did not display any increase in the distance between TM3-TM6 indicating that CCR5 stays in an inactive conformation (Fig. S19). Again, this behavior indicated the importance of considering protonated GLU283^7.39^ for GP120 bound state to observe activation process in MD simulations.

Isaikana et al. in their study suggested important activation switches for CCR5 receptor.^28^ They mentioned the relocalizations observed in TM6 and TM7 are due to changes in residues MET287^7.43^, HIS289^7.45^ and TRP248^6.48^. Zhang et al. who resolved the structure of CCR5 in complex with two natural chemokines and in apo state, indicate the importance of TYR251^6.51^ residue in addition to TM6 residue TRP248^6.48^ for activation process. Hence, we checked the orientations of these four residues in GP120 and MRV bound states by using selected structures from MD simulations (specifically structures with highest Δ*d* values were selected). We compared their orientations to the case where CCR5 was active and bound by super agonist [6P4]CCL5 as Isaikana et al. indicated that in this chemokine bound active state the orientations differ compared to inactive state. The TM6 residue TYR251^6.51^ adopted a similar orientation as in the active CCR5 structure when GP120 bound to CCR5 and the residues on both systems could be aligned on top of each other though not perfectly (Fig. 5a). Though for MRV bound state this residue could not be aligned to active state orientation which could indicate there was no movement in TM6 for MRV bound state (Fig. 5a). However, when we examined other TM6 residue TRP248^6.48^, we saw that in both GP120 bound and MRV bound states this residue adopts quite a different and dissimilar orientation than active state one (Fig. 5a and 5b). This residue was said to trigger the activation switches in CCR5 and one residue for relocalizations of TM6. For GP120 bound CCR5 the conformation of HIS289^7.45^ was closer to active one in an extended conformation, albeit side chains could not be aligned on top of each other (Fig. 5a). This alignment issue of GP120 bound CCR5 to active agonist bound state did not however indicated that GP120 bound state undergoes TM6 and TM7 relocalization, as mentioned before there were obvious rearrangements observed in intracellular TM sides for GP120 bound case. In fact, the Δ*d* value for For MRV bound CCR5 though HIS289^7.45^ had a different conformation as the side chain of the residue was not extended to the bundle of TMs as in the active conformation (Fig. 5a). The other TM7 residue MET287^7.43^ did not have the specified active state conformation for either of bound CCR5 systems (Fig. 5a and 5b). Isaikana et al. however indicated the importance of conformational changes in the side chain of this residue and how it brings notable local changes in the backbone of TM7. Hence, we also examined other active state structures resolved by Zhang et al.^32^ and observed that in fact MET287^7.43^ did not adopt a kind of downward orientation in chemokine bound states as we can see from the comparison of CCL3 bound and [6P4]CCL5 bound CCR5 structures (Fig. 5c). Hence, the conformational change in the side chain of this residue may not be necessary for receptor activation and could indicate a distinct activation mode as suggested by Zhang et al.^32^ with a different intermediate state of CCR5 receptor.

**Figure 5.**
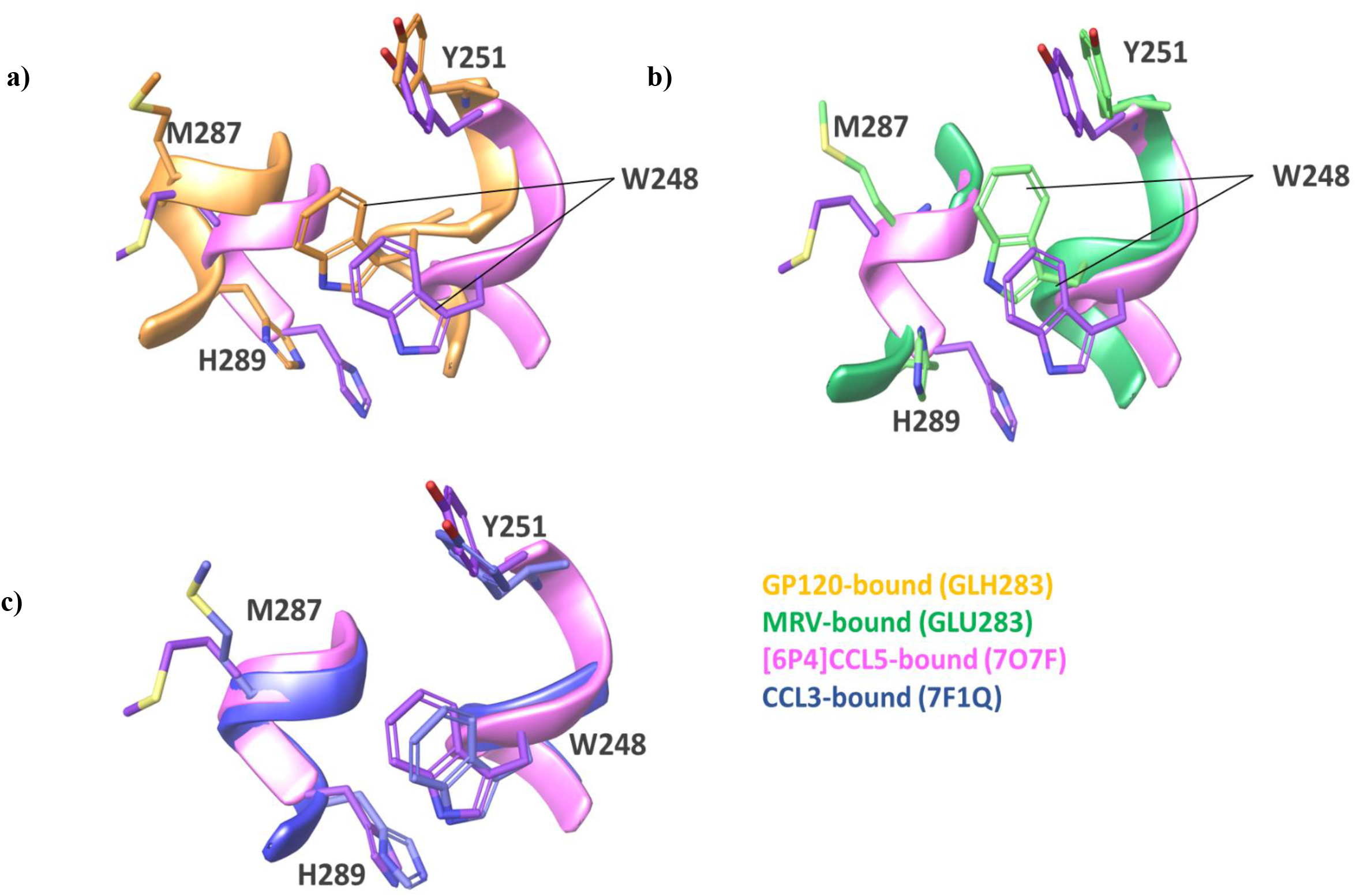
The comparasion of orientation for critical residues in TM6 and TM7 in CCR5 receptor bound by different ligands.

There were also conformational changes observed in the conserved motif for the activation process of class A GPCRs. In our case, we analyzed this change dynamically by calculating the RMSD of the Cα atoms in motifs for simulated systems relative to inactive structure (PDB ID, 4MBS). We compared the RMSD changes observed for GP120 bound CCR5 structure with protonated (GLH283^7.39^) system with MRV bound CCR5 structure with ionized (GLU283^7.39^) to observe whether GP120 binding could affect these switches differently than MRV binding could. The cut-off values to indicate significant changes were determined by calculating RMSD between resolved single point inactive and active structures (structures with PDB ID, 7F1S considered for active conformation). The first structural change we examined was in the conserved motif NPxxY which concurs with changes occurring in TM6 and TM7.^28, 69^ For CCR5, the corresponding residues in this motif were ASN293^7.49^, PRO294^7.50^, ILE295^7.51^, ILE296^7.52^ and TYR297^7.53^. The RMSD values larger than 2.85 Å were considered to indicate receptor adopting an active conformation. Based on the cut-off, we saw that there were a few MD frames which could be considered to represent active-like CCR5 structure with GP120 bound state though none of the collected structures for MRV bound state crossed the cut-off value for this motif (Fig. 6, top-left plot). Another motif we considered was PIF motif with corresponding residues of CCR5 being PRO206^5.50^, ILE116^3.40^ and TYR244^6.44^ and Isaikana et al.^28^ indicated to cause large-scale movements in TM6. For this motif, the cut-off RMSD value to indicate active conformation was taken as 1.57 Å. In this case, suprisingly we observed that for both MRV and GP120 bound states RMSD values larger than cut-off were observed. Though still GP120 bound CCR5 structure moved farther away from the inactive conformation based on the values (Fig. 6, top-right plot). DRY motif was another activation switch considered for class A GPCRs and intrahelical salt-bridge between ASP125^3.49^ and ARG126^3.50^ which is present in inactive state breaks druing activation process to open the binding pocket for G-protein.^66^ For this motif, the cut-off value was taken as 1.10 Å and we also checked whether salt-bridge interaction maintained (Fig. S17). We have seen that though for both GP120 and MRV bound CCR5 structures the RMSD values of the motif were higher than the cut-off at different times of MD simulations, it was GP120 bound state for which the values were larger and also salt-bridge interaction were mostly broken (only maintaned 23% of MD times) while this salt-bridge was more persistent for MRV bound state (maintaned 60% of MD times). The last motif we considered was TXP motif and it was the one suggested to be critical by Zhang et al.^31^ They monitored the RMSD changes of this motif in their accelarated MD simulations of wild type and mutated CCR5 structures to distinguish conformational states of the receptor. This motif is only found in chemokine receptor and a few peptide receptors^31^ and consists of residues TRP82^2.56^, VAL83^2.57^, PRO84^2.58^. The cut-off value we considered was 1.55 Å though it was taken as even a lower value by Zhang et al. In this case, we have seen that for GP120 bound state there were again a few MD frames which may be deemed as active conformation. All in all, these changes in RMSD of motifs as well as breakage of intrahelical salt-bridge in GP120 bound state taken together with distance analysis indicate CCR5 adopted a more active-like conformation with protonated GLU283^7.39^ residue.

**Figure 6.**
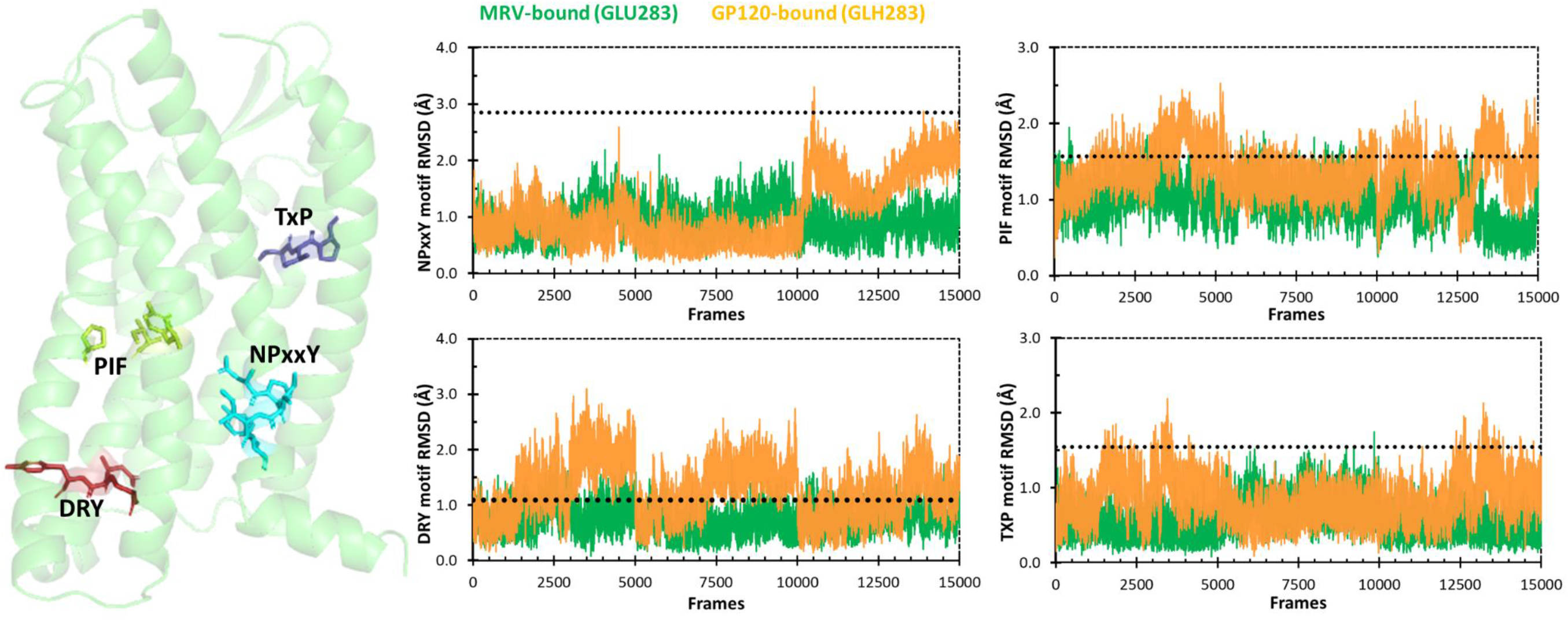
RMSD values for the conserved motifs in CCR5 for bound states relative to inactive CCR5 structure (PDB ID, 4MBS).The dotted black lines indicate the RMSD difference determined between inactive and active (PDB ID, 7F1S) crystal CCR5 structures.

## 4. CONCLUSIONS

In this study, we have performed MD simulations to investigate CCR5 coreceptor conformational heterogeneity. We utilized the structures of CCR5 obtained by cryo-EM in which it is in complex with GP120 as well as X-ray structure of CCR5 in complex with small molecule MRV. The replicate atomistic MD simulations were run with different protonation states of GLU283^7.39^, an important binding site residue in the cases of CCR5 bound by either GP120 or MRV as well as in *apo* state. The various post-MD analysis we performed showed that the protonation state of GLU283^7.39^ caused differences in conformational state of CCR5 in terms of its adopting active or inactive-like conformation. Especially in the case of GP120 bound CCR5 with protonated GLU283^7.39^ residue we observed that the coreceptor can assume an active-like state that is different that the case with ionized GLU283^7.39^ case. The conformation adopted was also distint to the one adopted by MRV bound CCR5 as well as the *apo* form conformation. This observation can have some insinuations especially for novel inhibitor molecules against CCR5 using approaches such as molecular docking as in many of the studies only the MRV bound conformation of this coreceptor with ionized GLU283^7.39^ was considered. However, considering the CCR5 conformation that is adopted when it is bound by GP120 could lead to discovery of compounds with higher binding potential and would be part of our future work.

## ACKNOWLEDGMENTS

The numerical calculations reported in this paper were fully performed at TUBITAK ULAKBIM, High Performance and Grid Computing Center (TRUBA resources). This study has been financially funded by Scientific and Technological Research Council of Turkey (TUBITAK) under the program of TUBITAK-2218. BD also gratefully acknowledges funding by Istanbul Technical University BAP THD-2023-44790 project.

The authors declare that there is no conflict of interest regarding the publication of this article.

