## Supplementary Material for "Investigating the Effect of GLU283 Protonation State on the Conformational Heterogeneity of CCR5 by Molecular Dynamics Simulations"

**Table S1.** Publicly available structures of CCR5 coreceptor.

| PDB ID | State | Ligand | Year resolved | Method used |
|---|---|---|---|---|
| 4MBS | inactive | Maraviroc (antagonist) | 2013 | X-ray |
| 5UIW | inactive | 5P7-CCL5 (antagonist) | 2017 | X-ray |
| 6AKX | inactive | Compound 21 (antagonist) | 2018 | X-ray |
| 6AKY | inactive | Compound 34 (antagonist) | 2018 | X-ray |
| 6MEO | inactive | HIV-1 envelope protein GP120 | 2018 | Cryo-EM |
| 7OF7 | active | [6P4]CCL5 (agonist) and $G_i$ | 2021 | Cryo-EM |
| 7F1Q | active | MIP-1α (agonist) and $G_i$ | 2021 | Cryo-EM |
| 7F1R | active | RANTES (agonist) and $G_i$ | 2021 | Cryo-EM |
| 7F1S | active | No ligand, $G_i$ only | 2021 | Cryo-EM |
| 7F1T | inactive | MIP-1α (agonist) | 2021 | X-ray |

**Table S2.** The conditions of replica molecular dynamics simulations.

| State | Simulation ID | Starting PDB ID | p$K_a$ of GLU283[ǂ] | GLU283 | Duration of MD simulation |
|---|---|---|---|---|---|
| *CD4/GP120 bound* | MD-1 | 6MEO | 8.38 | ionized | 3 x 500 ns = 1.5 µs |
| | MD-2 | 6MEO | 8.38 | protonated | 3 x 500 ns = 1.5 µs |
| *HOLO* | MD-3 | 4MBS | 5.48 | ionized | 3 x 500 ns = 1.5 µs |
| | MD-4 | 4MBS | 5.48 | protonated | 3 x 500 ns = 1.5 µs |
| *APO* | MD-5 | 7F1S | 7.11 | ionized | 3 x 500 ns = 1.5 µs |
| | MD-6 | 7F1S | 7.11 | protonated | 3 x 500 ns = 1.5 µs |
| | MD-7 | 6MEO | 6.19 | ionized | 3 x 500 ns = 1.5 µs |
| | MD-8 | 6MEO | 6.19 | protonated | 3 x 500 ns = 1.5 µs |
| | MD-9 | 4MBS | 5.76 | ionized | 3 x 500 ns = 1.5 µs |
| | MD-10 | 4MBS | 5.76 | protonated | 3 x 500 ns = 1.5 µs |

*Ligand molecule in MD-3 and MD-4 is Maraviroc and when GLU283 is protonated, the nitrogen atom of tropane ring is negatively charged

[ǂ]Calculated using PROPKA module implemented in Schrodinger simulation package.

a)

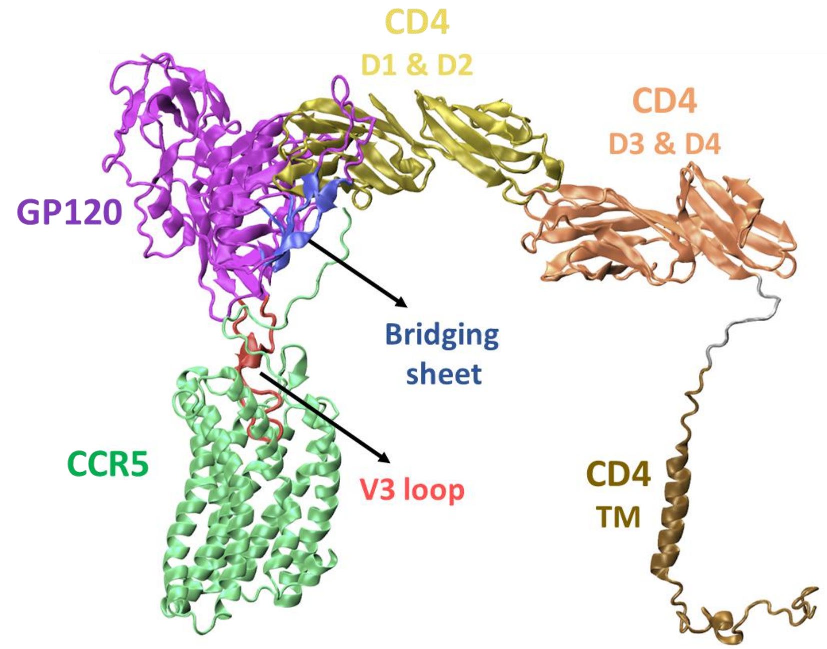


b)

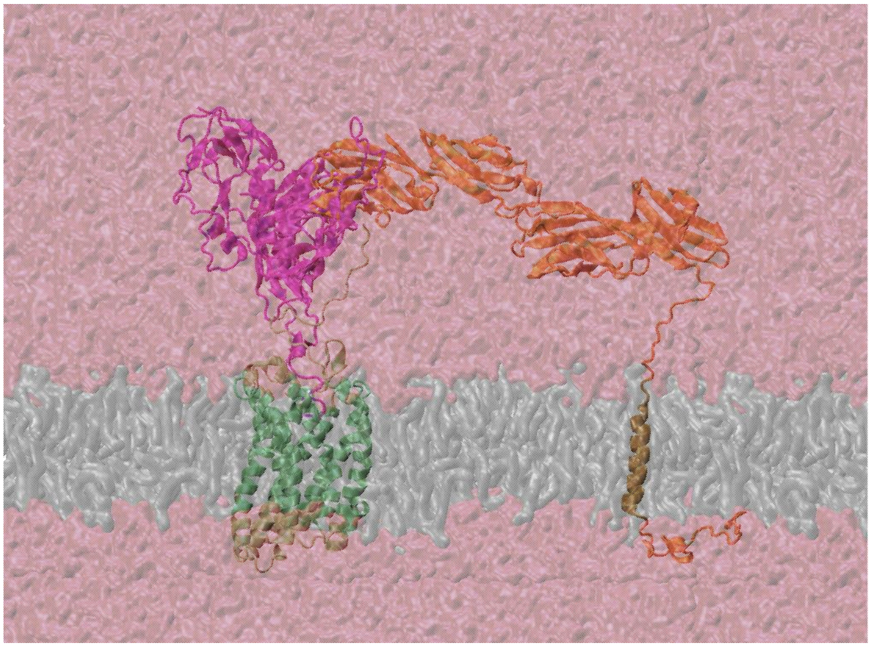

**Figure S1**. Simulation system setups for CD4-GP120 bound CCR5 a) ribbon representation with domains of CD4 receptor are shown in different colors and bridging sheet, as well as V3 loop of GP120, are displayed in blue and red colors respectively. b) simulation box with periodic boundary conditions with membrane layer represented as gray transparent surface and water/ion layer represented as red transparent surface.

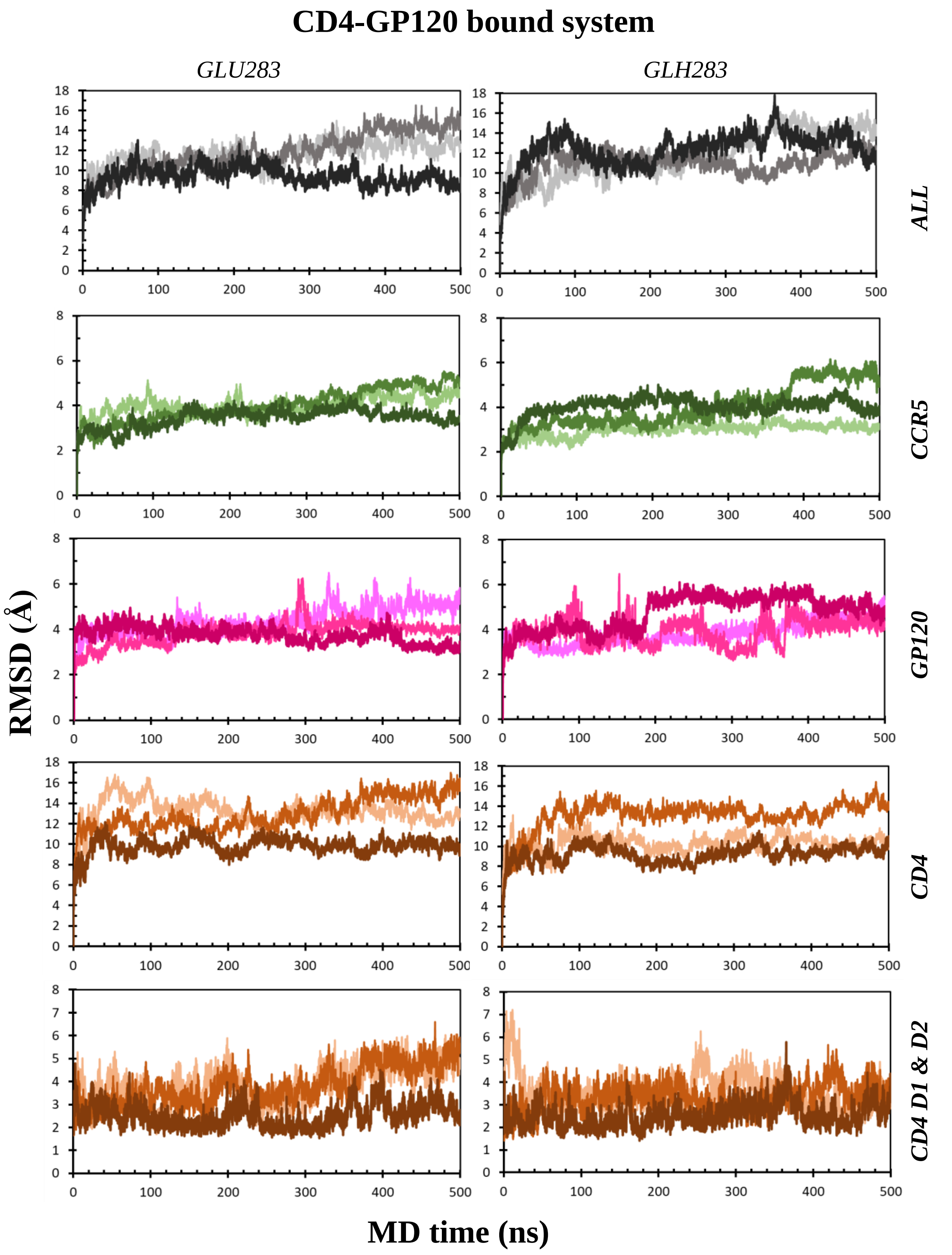


**Figure S2**. Root mean square deviation (RMSD) values of Cα atoms observed during MD simulation times for *CD4-/GP120* bound CCR5 systems, i.e. for three replicate simulations with simulations IDs MD-1 and MD-2 in Table S1.

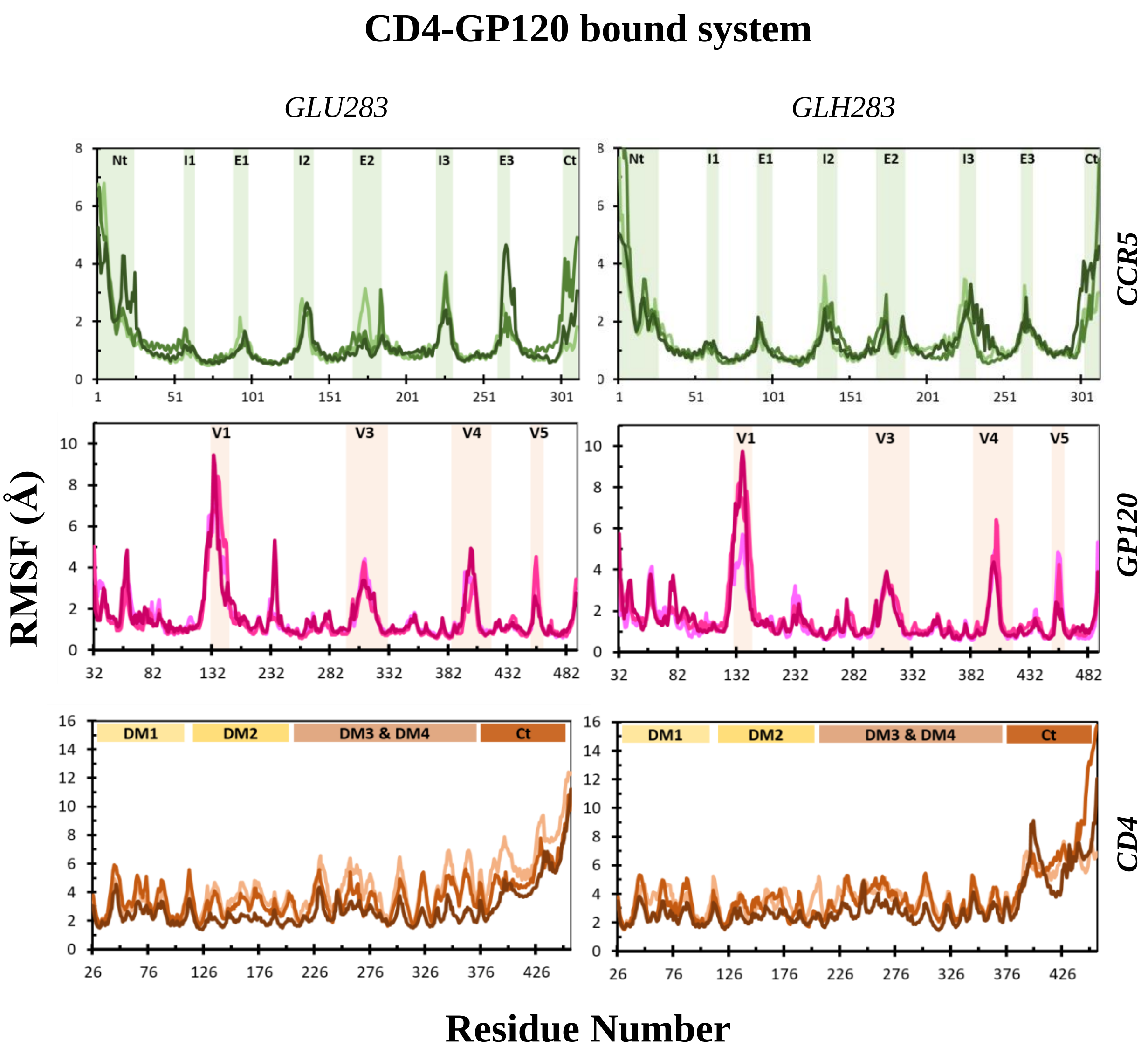


**Figure S3.** Root mean square fluctuations (RMSF) of the Cα atoms of protein systems (for three replicate simulations with simulations IDs MD-1 and MD-2 in Table S1). Colored areas in graphs display the different regions of proteins. For CCR5 co-receptor, Nt represents N-terminus region, I1 represents intracellular loop 1, E1 represents extracellular loop 1, I2 represents intracellular loop 2, E2 represents extracellular loop 2, I3 represents intracellular loop 3, E3 represents extracellular loop 3 and Ct represents C-terminus. For GP120 glycoprotein, V1 represents variable loop 1, V3 represents variable loop 3, V4 represents variable loop 4 and V5 represents variable loop 5. For CD4 receptor, DM1 represents the domain 1, DM2 represents the domain 2 and DM3 & DM4 represents the domains 3 and 4 respectively (See figure S1 for domains of CD4 receptor).

### MRV-bound system

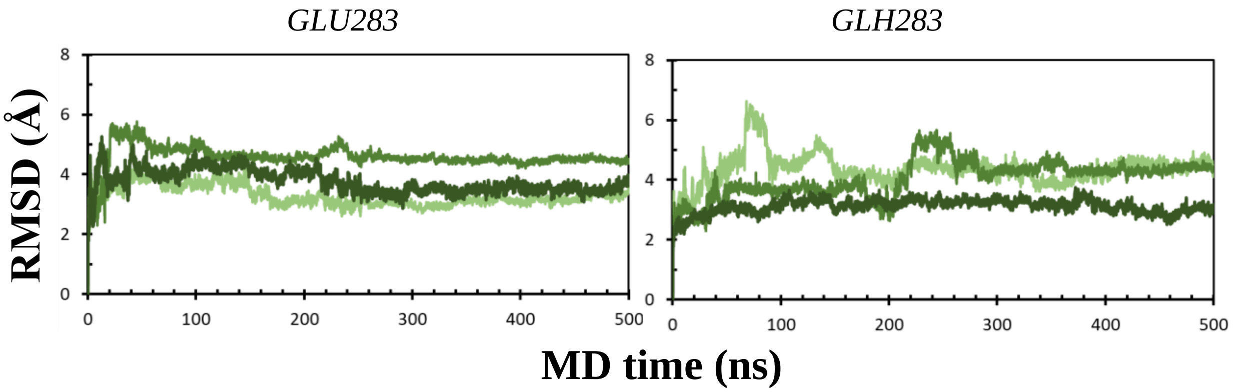


**Figure S4**. RMSD values of Cα atoms observed during MD simulation times for *MRV*-bound CCR5 systems, i.e. for three replicate simulations with simulations IDs MD-3 and MD-4 in Table S1.

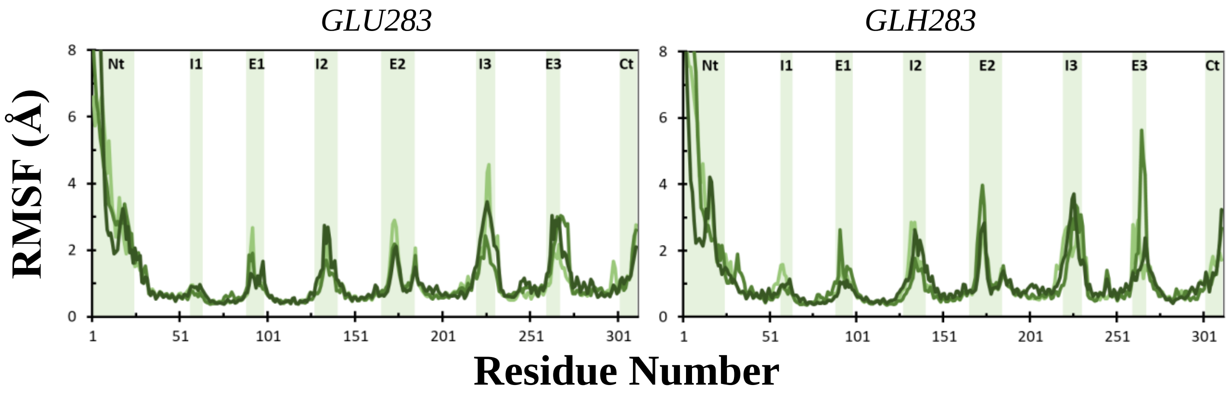


**Figure S5.** RMSF of the Cα atoms of protein systems (for three replicate simulations with simulations IDs MD-3 and MD-4 in Table S1). Colored areas in graphs display the different regions of proteins. For CCR5 co-receptor, Nt represents N-terminus region, I1 represents intracellular loop 1, E1 represents extracellular loop 1, I2 represents intracellular loop 2, E2 represents extracellular loop 2, I3 represents intracellular loop 3, E3 represents extracellular loop 3 and Ct represents C-terminus.

### APO system – initiated from active structure (PDB ID: 7F1S)

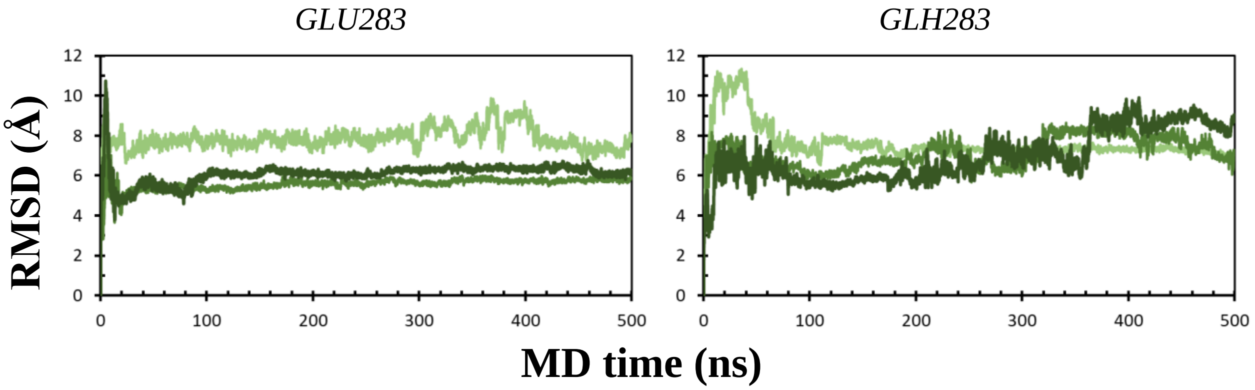


**Figure S6**. RMSD values of Cα atoms observed during MD simulation times for *apo* CCR5 systems, i.e. for three replicate simulations with simulations IDs MD-5 and MD-6 in Table S1.

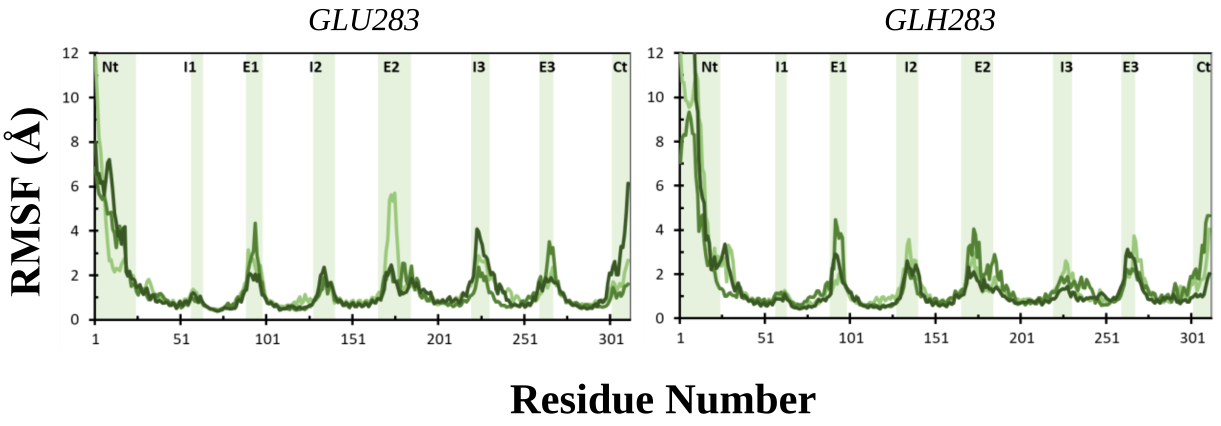


**Figure S7.** RMSF of the Cα atoms of protein systems (for three replicate simulations with simulations IDs MD-5 and MD-6 in Table S1). Colored areas in graphs display the different regions of proteins. For CCR5 co-receptor, Nt represents N-terminus region, I1 represents intracellular loop 1, E1 represents extracellular loop 1, I2 represents intracellular loop 2, E2 represents extracellular loop 2, I3 represents intracellular loop 3, E3 represents extracellular loop 3 and Ct represents C-terminus.

### APO system – initiated from inactive structure (PDB ID: 6MEO)

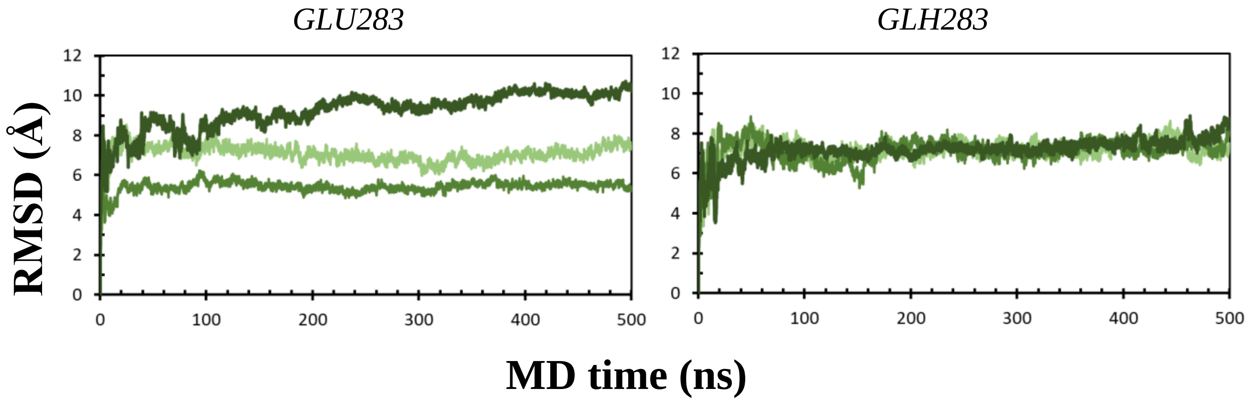


**Figure S8**. RMSD values of Cα atoms observed during MD simulation times for *apo* CCR5 systems, i.e. for three replicate simulations with simulations IDs MD-9 and MD-10 in Table S1.

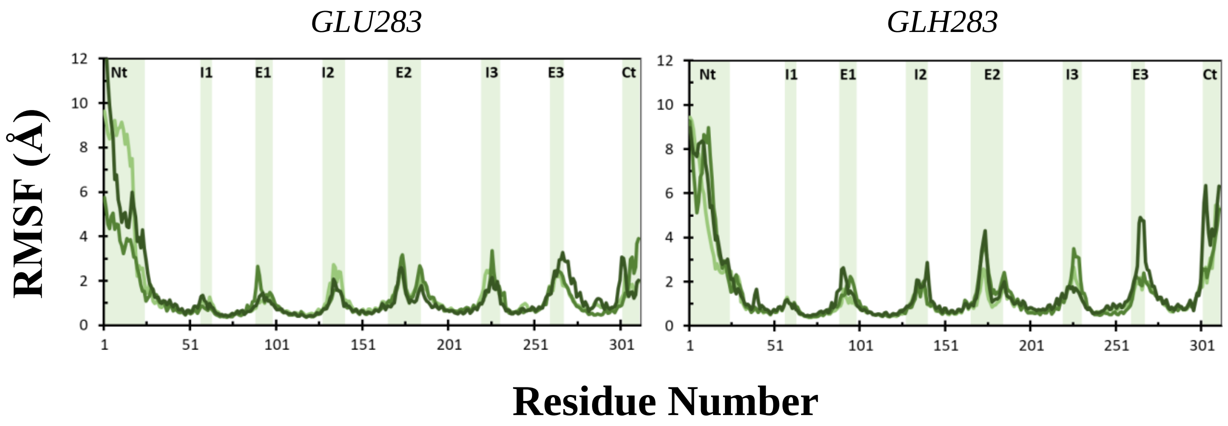


**Figure S9.** RMSF of the Cα atoms of protein systems (for three replicate simulations with simulations IDs MD-9 and MD-10 in Table S1). Colored areas in graphs display the different regions of proteins. For CCR5 co-receptor, Nt represents N-terminus region, I1 represents intracellular loop 1, E1 represents extracellular loop 1, I2 represents intracellular loop 2, E2 represents extracellular loop 2, I3 represents intracellular loop 3, E3 represents extracellular loop 3 and Ct represents C-terminus.

### APO system – initiated from inactive structure (PDB ID: 4MBS)

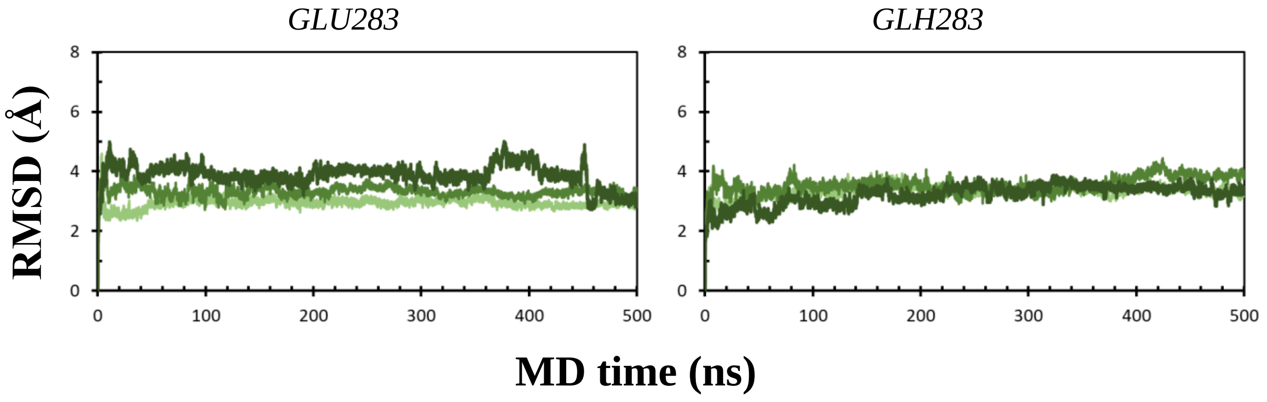


**Figure S10**. RMSD values of Cα atoms observed during MD simulation times for *apo* CCR5 systems, i.e. for three replicate simulations with simulations IDs MD-7 and MD-8 in Table S1.

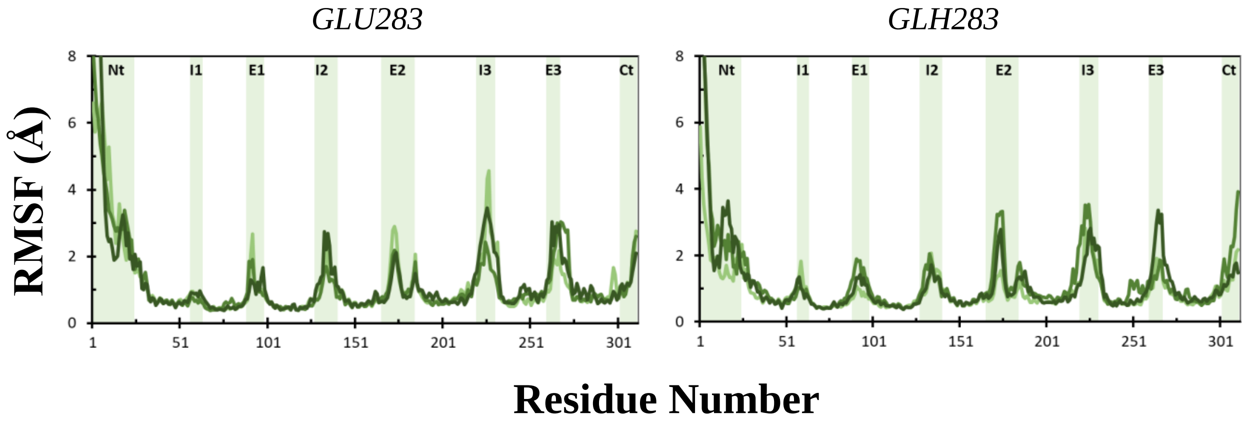


**Figure S11.** RMSF of the Cα atoms of protein systems (for three replicate simulations with simulations IDs MD-7 and MD-8 in Table S1). Colored areas in graphs display the different regions of proteins. For CCR5 co-receptor, Nt represents N-terminus region, I1 represents intracellular loop 1, E1 represents extracellular loop 1, I2 represents intracellular loop 2, E2 represents extracellular loop 2, I3 represents intracellular loop 3, E3 represents extracellular loop 3 and Ct represents C-terminus.

| Interaction Type | GP120 residue | CCR5 residue | % Frequency (GLU283) | % Frequency (GLH283) |
|---|---|---|---|---|
| Salt bridges | ARG298 | TYR14 | 35 | 39 |
| | ARG304 | GLU172 | 93 | 99 |
| | ARG313 | ASP276 | 1 | 62 |
| | ARG313 | GLU(H)283 | 39 | 0 |
| | ASP320 | LYS22 | 69 | 35 |
| | ASP320 | LYS26 | 19 | 44 |
| | LYS416 | TYR10 | 31 | 35 |
| | LYS435 | TYR14 | 13 | 25 |
| Hydrogen bonds | ARG298 | TYR14 | 53 | 58 |
| | ASN302 | TYR14 | 85 | 90 |
| | ARG304 | GLU172 | 93 | 99 |
| | SER306 | SER179 | 55 | 46 |
| | ILE307 | SER180 | 97 | 91 |
| | HIS308 | TYR89 | 66 | 24 |
| | HIS308 | CYS178 | 2 | 48 |
| | GLY312 | GLU(H)283 | 99 | 26 |
| | ARG313 | GLU(H)283 | 82 | 0 |
| | ARG313 | ASN258 | 11 | 63 |
| | ARG313 | ASP276 | 1 | 61 |
| | ALA314 | GLN280 | 23 | 52 |
| | TYR316 | TYR89 | 50 | 54 |
| | ASP320 | LYS22 | 68 | 36 |
| | ASP320 | LYS26 | 19 | 43 |
| | ARG326 | ASP11 | 74 | 98 |
| | LYS416 | TYR10 | 76 | 77 |
| | GLN417 | TYR10 | 99 | 89 |
| | ILE418 | TYR10 | 83 | 64 |
| | PRO433 | TYR15 | 100 | 99 |
| | LYS435 | TYR14 | 44 | 69 |
| | GLY436 | TYR14 | 65 | 52 |
| $\pi$-cation | LYS207 | TYR15 | 17 | 33 |
| | ARG326 | TYR10 | 59 | 46 |
| | ARG414 | TYR10 | 40 | 47 |
| | LYS435 | TYR14 | 49 | 73 |

**Figure S12**. Comparison of interactions between GP120 and CCR5 with ionized GLU283 and protonated GLH283. Replica MD simulations were concatenated for both systems and used for the dynamical evaluation of interactions. The persistency of interactions is given in percentages, representing how much of the simulation time they are maintained.

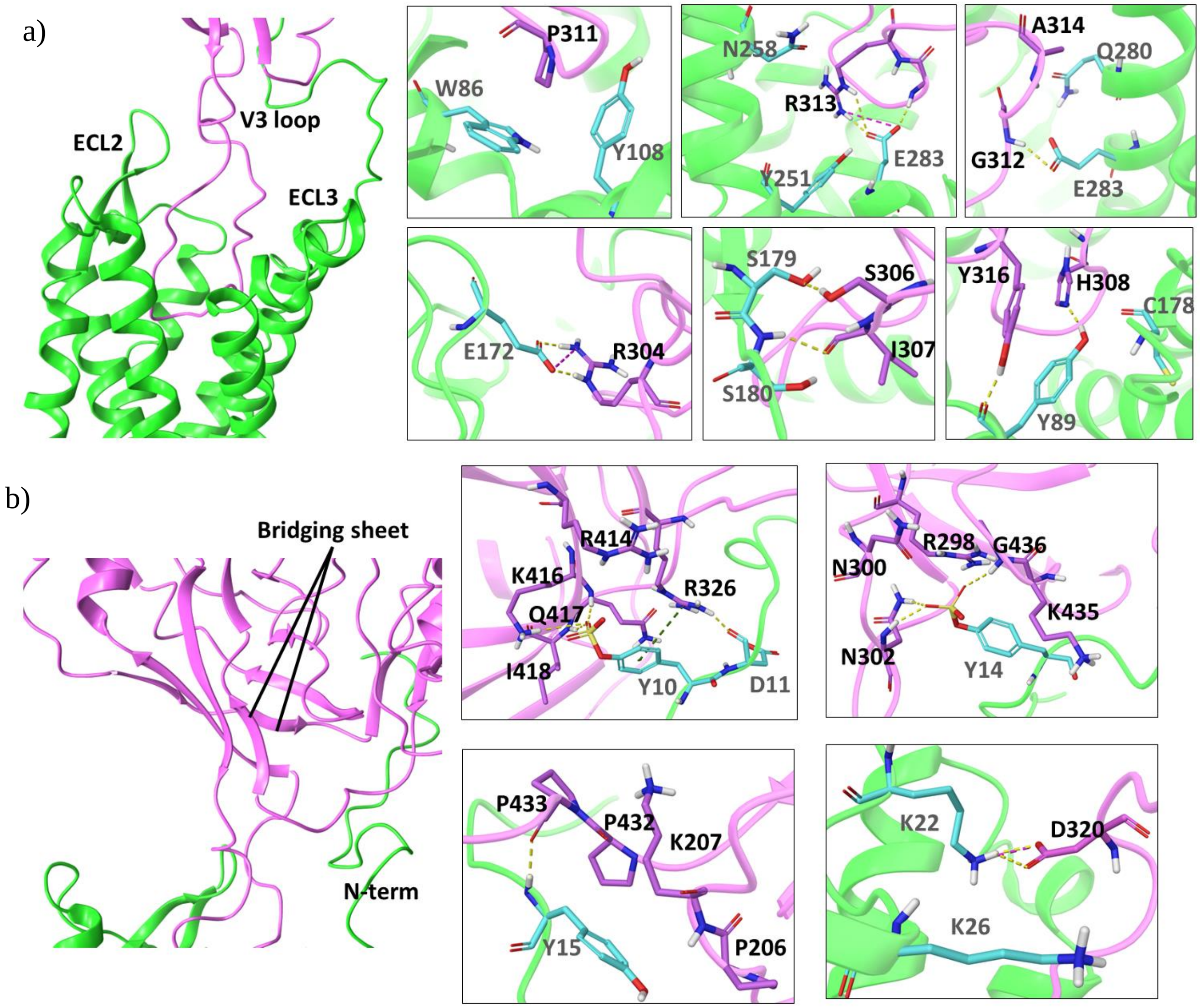


**Figure S13**. The interactions between GP120 and CCR5 with ionized GLU283[7.39]. a) for V3 loop of GP120 and chemokine binding site 2 of CCR5 b) for bridging sheet of GP120 and N-term of CCR5. Dashed yellow lines represent hydrogen bonds; green ones represent pi-pi; pink ones represent salt-bridge interactions.

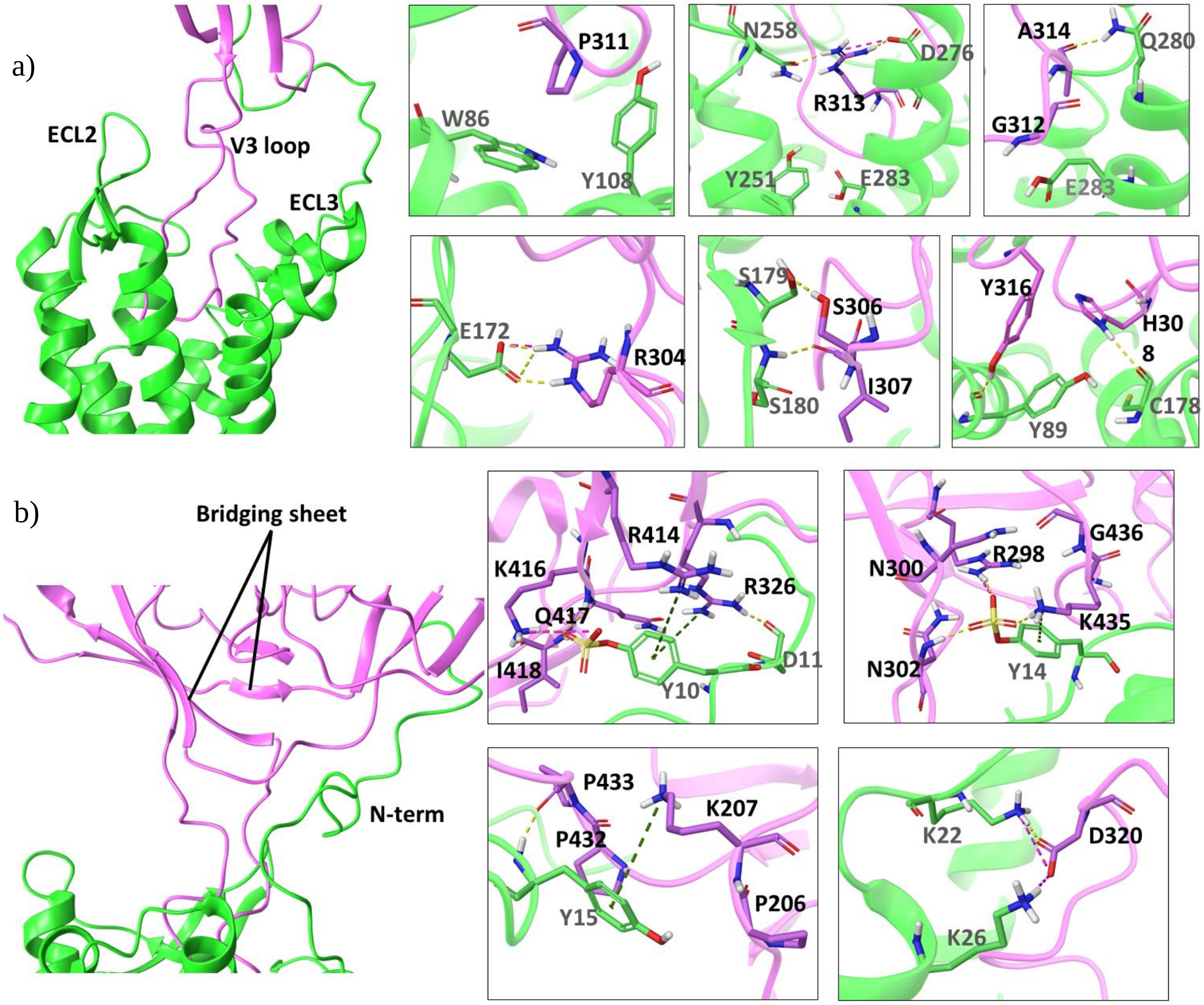


**Figure S14**. The interactions between GP120 and CCR5 with protonated GLH283[7.39] a) for V3 loop of GP120 and chemokine binding site 2 of CCR5 b) for bridging sheet of GP120 and N-term of CCR5. Dashed yellow lines represent hydrogen bonds; green ones represent pi-pi; pink ones represent salt-bridge interactions.

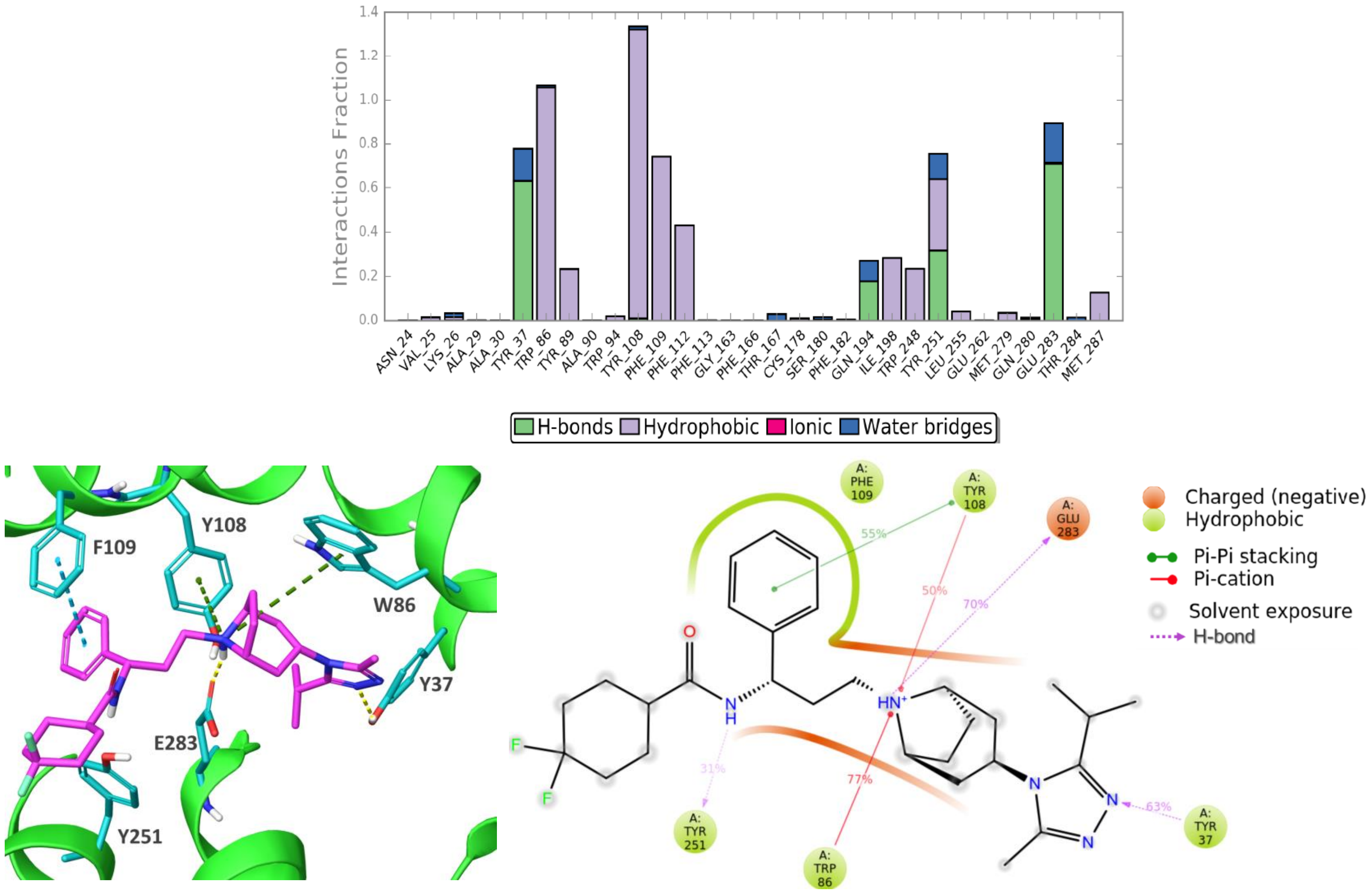


**Figure S15**. The interactions between MRV and CCR5 with ionized GLU283[7,39]

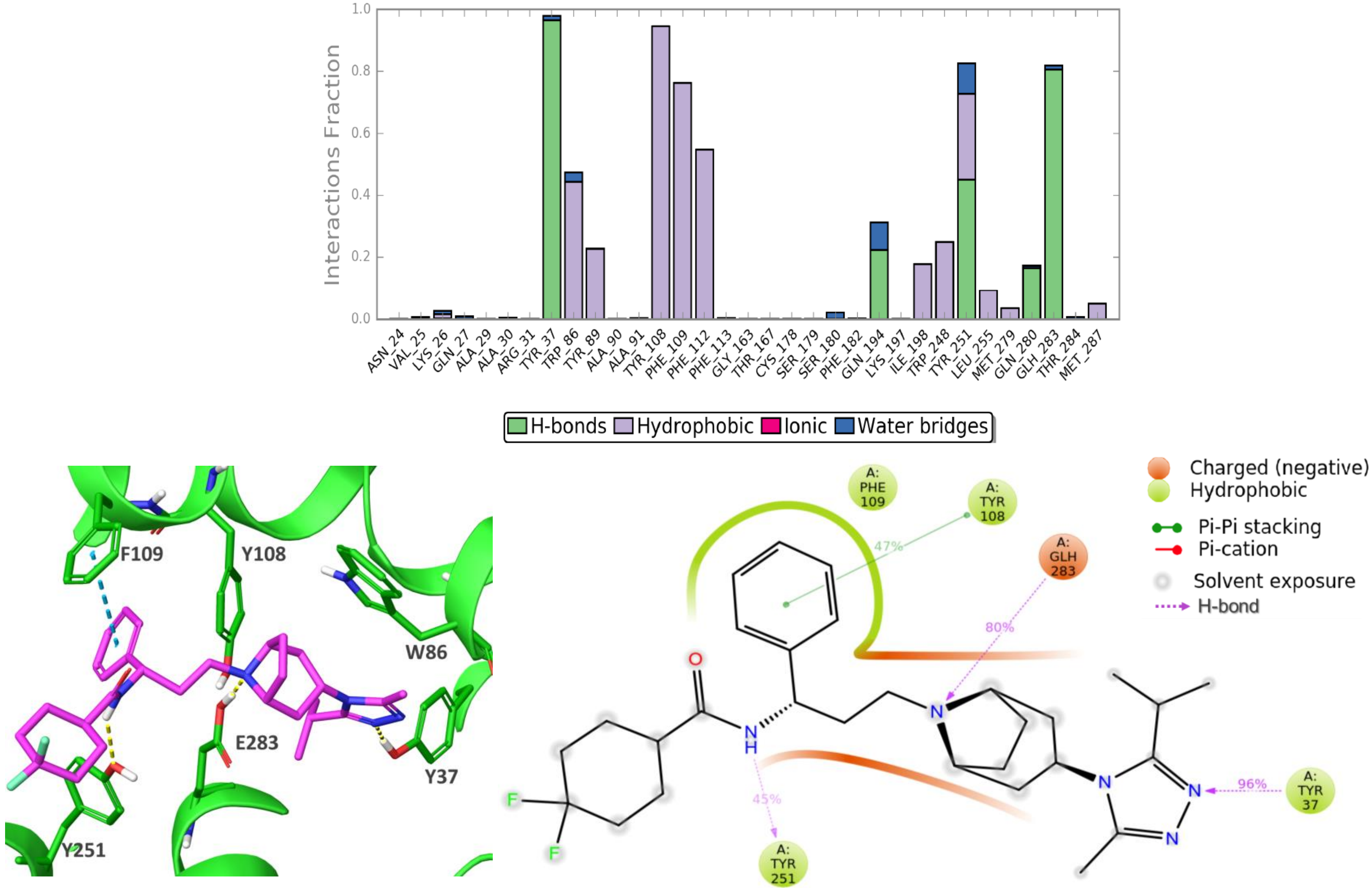


**Figure S16.** The interactions between MRV and CCR5 with protonated GLH283[7,39]

| | Residue A | Residue B | GP120 | | HOLO | | APO | | | | | |
|---|---|---|---|---|---|---|---|---|---|---|---|---|
| | | | (GLU) | (GLH) | (GLU) | (GLH) | 7F1S (GLU) | 7F1S (GLH) | 6MEO (GLU) | 6MEO (GLH) | 4MBS (GLU) | 4MBS (GLH) |
| **Salt bridges** | ASP66 | LYS59 | 98 | 90 | 99 | 98 | 98 | 99 | 93 | 88 | 90 | 99 |
| | ASP125 | ARG126 | 7 | 23 | 60 | 52 | 15 | 43 | 37 | 68 | 40 | 14 |
| | ASP125 | ARG140 | 100 | 97 | 100 | 99 | 100 | 65 | 95 | 81 | 100 | 100 |
| | ARG230 | GLU227 | 78 | 85 | 1 | 0 | 66 | 89 | 95 | 88 | 0 | 0 |
| | ARG232 | GLU302 | 37 | 39 | 60 | 64 | 34 | 34 | 51 | 72 | 61 | 95 |
| | GLU262 | LYS191 | 32 | 33 | 85 | 51 | 23 | 44 | 0 | 0 | 67 | 77 |
| | ASP276 | LYS22 | 30 | 34 | 57 | 86 | 86 | 88 | 44 | 14 | 30 | 79 |
| **Hydrogen bonds** | ASH76 | ASN48 | 7 | 72 | 95 | 100 | 80 | 82 | 25 | 19 | 64 | 95 |
| | ASH76 | ASN293 | 76 | 72 | 72 | 70 | 85 | 40 | 35 | 28 | 73 | 95 |
| | ASH76 | TYR297 | 64 | 13 | 7 | 0 | 0 | 0 | 7 | 0 | 5 | 1 |
| | PRO84 | TYR89 | 0 | 0 | 0 | 0 | 64 | 88 | 0 | 0 | 0 | 0 |
| | TYR108 | TYR251 | 36 | 17 | 16 | 0 | 69 | 93 | 24 | 54 | 37 | 68 |
| | TYR108 | GLU283 | 0 | 57 | 83 | 69 | 82 | 29 | 62 | 32 | 86 | 15 |
| | THR123 | TYR214 | 59 | 66 | 34 | 28 | 2 | 13 | 85 | 94 | 56 | 0 |
| | ASP125 | THR65 | 42 | 66 | 77 | 91 | 63 | 81 | 75 | 84 | 94 | 90 |
| | ASP125 | ARG140 | 100 | 97 | 100 | 99 | 100 | 65 | 95 | 81 | 100 | 100 |
| | GLN170 | THR177 | 91 | 67 | 99 | 95 | 33 | 4 | 99 | 97 | 89 | 65 |
| | CYS178 | TYR89 | 82 | 83 | 48 | 90 | 0 | 0 | 76 | 43 | 81 | 79 |
| | ARG232 | GLU302 | 36 | 39 | 58 | 64 | 33 | 34 | 49 | 80 | 61 | 95 |
| | TYR244 | ASN293 | 31 | 36 | 32 | 0 | 0 | 0 | 37 | 69 | 0 | 0 |
| | TYR244 | GLY202 | 0 | 0 | 18 | 75 | 89 | 73 | 33 | 23 | 82 | 92 |
| | TYR244 | ALA249 | 11 | 0 | 11 | 1 | 37 | 97 | 33 | 1 | 5 | 4 |
| | HIE289 | TRP248 | 54 | 80 | 60 | 68 | 82 | 65 | 57 | 87 | 96 | 97 |
| | ILE295 | VAL300 | 0 | 0 | 0 | 0 | 78 | 50 | 0 | 0 | 0 | 0 |
| | PHE299 | PRO294 | 0 | 0 | 0 | 0 | 70 | 49 | 0 | 0 | 0 | 0 |
| | ARG305 | PHE299 | 0 | 0 | 61 | 67 | 15 | 7 | 0 | 0 | 86 | 52 |
| **$\pi$-cation $\pi$-$\pi$** | ARG168 | HIS181 | 47 | 65 | 67 | 53 | 49 | 17 | 46 | 74 | 66 | 13 |
| | HIS289 | PHE247 | 25 | 20 | 9 | 7 | 36 | 72 | 10 | 3 | 23 | 10 |
| | HIS289 | TYR244 | 97 | 93 | 75 | 31 | 32 | 10 | 48 | 74 | 6 | 2 |
| | HIS289 | PHE79 | 40 | 55 | 51 | 49 | 0 | 1 | 44 | 71 | 2 | 1 |

**Figure S17**. Intra-receptor residue-residue interactions within CCR5 for all simulated systems.

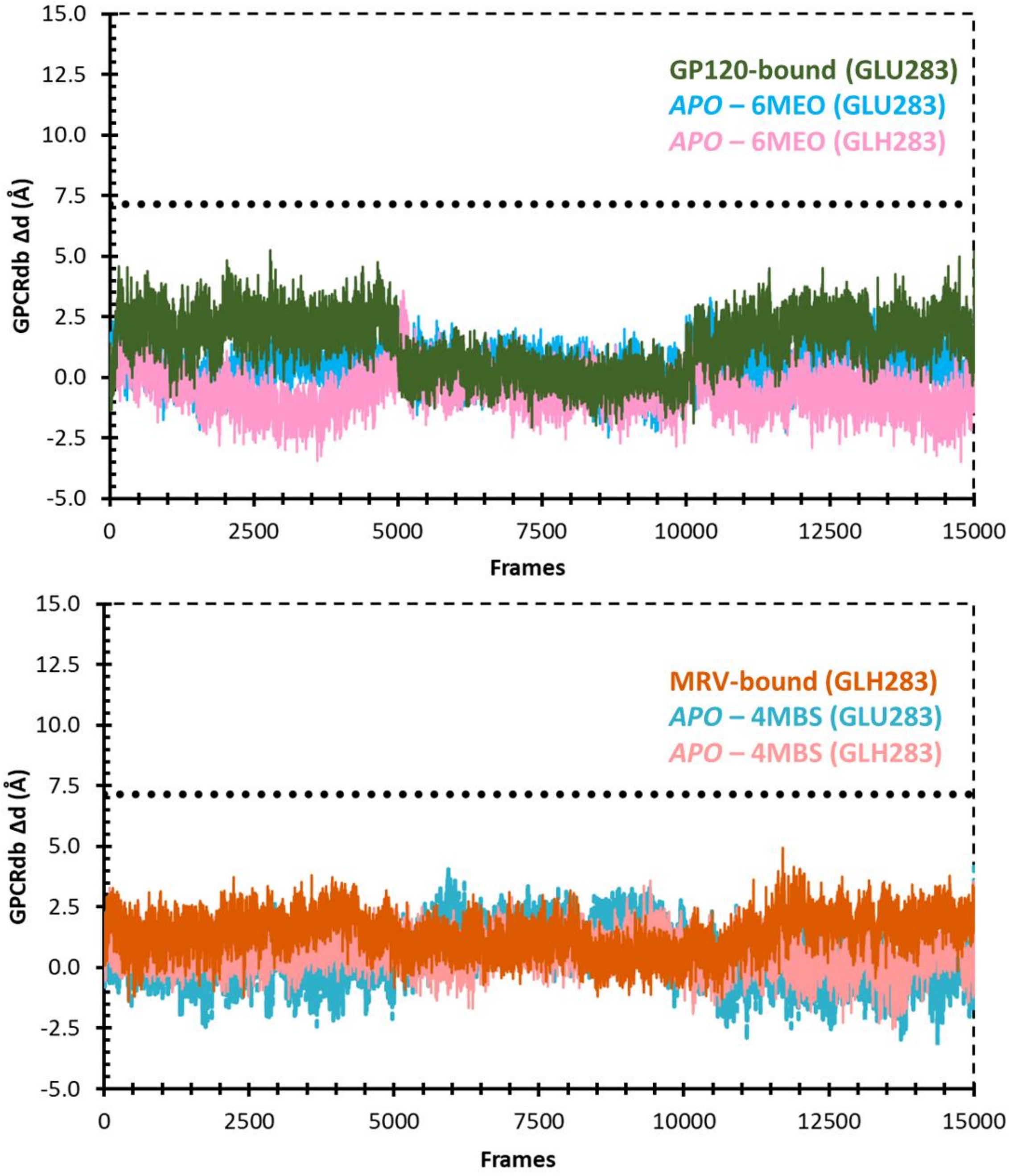


**Figure S18.** Δ*d* distance measured for the systems using concatenated MD trajectories

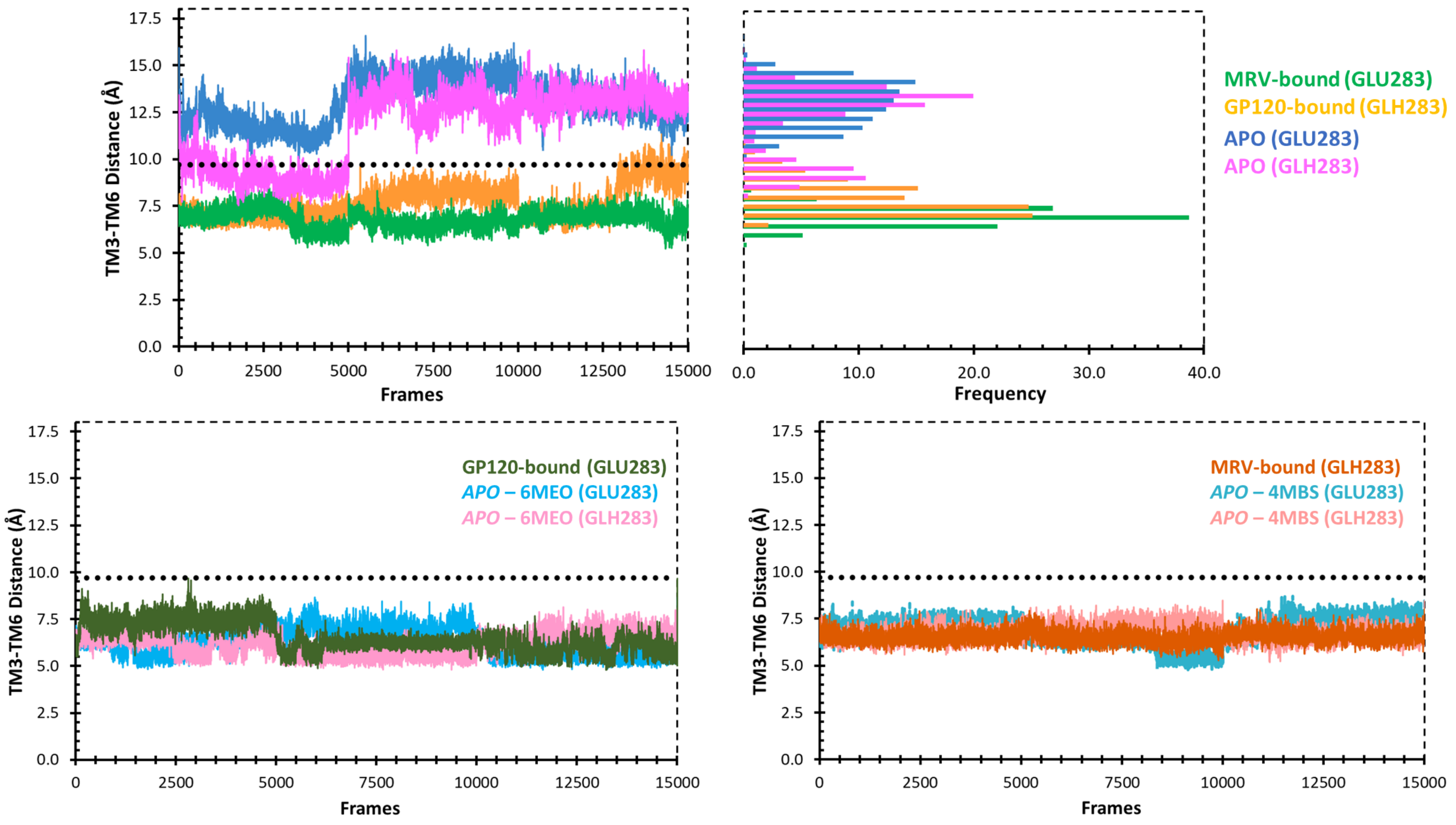


**Figure S19.** TM3-TM6 distance measured by calculating the distance between Cα of residues ARG126$^{3.50}$ and VAL234$^{6.34}$
